# MAP4K1 and MAP4K2 regulate ABA-induced and Ca^2+^-mediated stomatal closure in Arabidopsis

**DOI:** 10.1101/2024.07.05.602132

**Authors:** Kota Yamashita, Sotaro Katagiri, Hinano Takase, Yangdan Li, Airi Otoguro, Yoshiaki Kamiyama, Shota Yamauchi, Atsushi Takemiya, Izumi C. Mori, Taishi Umezawa

## Abstract

Abscisic acid (ABA)-induced stomatal closure limits water loss from plants under drought stress. To investigate signaling pathways involved in stomatal closure, we performed a phosphoproteomic analysis of ABA-treated Arabidopsis guard cell protoplasts (GCPs). From this analysis, we discovered that MITOGEN-ACTIVATED PROTEIN 4 KINASE 1 (MAP4K1) is phosphorylated in response to ABA. Subsequent biochemical assays showed that Ser-479 of MAP4K1 is directly phosphorylated by SRK2E (OST1/SnRK2.6), a protein kinase that plays a central role in ABA-induced responses. Mutational analyses of *MAP4K1*, as well as closely related *MAP4K2*, revealed that both kinases positively regulate ABA-induced stomatal closure, and that Ser-479 of MAP4K1 was required for this phenotype. In *map4k1map4k*2, stomatal closure was induced by exogenous Ca^2+^ but not H_2_O_2_. Furthermore, electrophysiology experiments showed that MAP4K1/2 are required for ABA-dependent activation of Ca^2+^-permeable channels in GCPs. Together, our results indicate that SnRK2 and MAP4K1/2 function as a signaling module to regulate Ca^2+^-mediated stomatal closure.

## Introduction

Stomata are small pores on the leaf epidermis that mediate gas exchange with the surrounding atmosphere. Each stoma consisting of a pair of guard cells surrounding a central pore. Stomatal aperture is tightly controlled in response to environmental changes. Stomata open in response to light and CO_2_ to promote gas exchange and transpiration for photosynthetic carbon assimilation and maintenance of leaf temperature. Conversely, stomata close to prevent water loss when plants experience drought or osmotic stress. The regulatory pathway of stomatal movements is crucial for plant growth and stress responses.

Stomatal closure is largely dependent on a phytohormone abscisic acid (ABA), and subclass III SNF1-related protein kinase 2 (SnRK2) kinases play a crucial role in ABA signaling. SRK2E (OST1/SnRK2.6) is a member of subclass III and plays a primary role in ABA-induced stomatal closure ^2,3^. ABA induces the activation of SRK2E, which directly phosphorylates SLOW ANION CHANNEL-ASSOCIATED1 (SLAC1), thereby promoting anion efflux ^4,5^. On the other hand, ABA induces an increase in reactive oxygen species (ROS) and cytoplasmic Ca^2+^ concentration ([Ca^2+^]_cyt_) in guard cells ^1^. In addition to activation of SRK2E, ABA-dependent [Ca^2+^]_cyt_ elevation activates several Ca^2+^-dependent protein kinases (CDPKs) and a pair of calcineurin-B-like proteins (CBLs) and CBL-interacting protein kinases (CIPKs) ^4,5^ to phosphorylate SLAC1 or SLAC1 homolog 3 (SLAH3) ^4,5^. These processes result in plasma membrane (PM) depolarization and a reduction in guard cell volume, leading to stomatal closure ^1^.Therefore, both Ca^2+^-dependent and -independent pathways coordinate ABA-induced stomatal closure ^4,5^.

ROS induce the activation of plasma membrane Ca^2+^-permeable (*I_Ca_*) channels ^6–8^.SRK2E is involved in this process based on several lines of evidence. ABA-dependent ROS production and [Ca^2+^]_cyt_ elevation are nearly abolished in a *srk2e/ost1* mutant, and SRK2E phosphorylates the membrane-bound and ROS-producing NADPH oxidases RESPIRATORY BURST OXIDASE HOMOLOG F and D (RBOHF/D) to promote ROS production ^9–13^.These findings suggest a connection between SRK2E and the Ca^2+^-dependent pathway, though a mechanistic explanation for this connection remains unknown.

In this study we identified mitogen-activated protein kinase kinase kinase kinase 1 (MAP4K1/AtMAP4Kα1) and MAP4K2 as positive regulators of ABA-induced stomatal closure. Our results further demonstrate that SRK2E directly phosphorylates MAP4K1 at Ser-479, and that MAP4K1/2 mediates ABA signaling to elevate [Ca^2+^]_cyt_. Taken together, our results significantly expand current models of ABA-induced stomatal closure.

## Results

### An overview of phosphoproteomic analysis using Arabidopsis guard cell protoplasts

Previous phosphoproteomic studies of ABA-induced responses in Arabidopsis seedlings have identified multiple SnRK2 substrates ^14–16^. In this study, we carried out a cell type-specific phosphoproteomic analysis to identify SnRK2 substrates in guard cells. Using a previously developed protocol^17,18^, guard cell protoplasts (GCPs) were isolated from leaves of Arabidopsis wild type (Col-0) and two types of ABA-insensitive mutants: *srk2de*, a double knockout mutant of SRK2D (SnRK2.2) and SRK2E (OST1/SnRK2.6), and *abi1-1C*, a gain-of-function mutant of type 2C protein phosphatase ABA INSENSITIVE 1 (ABI1)^19^. Prior to phosphoproteomic analysis, a far-western blot analysis was performed to confirm ABA-induced phosphorylation of ABA-RESPONSIVE KINASE SUBSTRATE (AKS) transcription factors ^20,21^. As reported previously, ABA-responsive AKSs were phosphorylated in Col-0 GCPs, and levels of AKS phosphorylation were reduced in *srk2de* and *abi1-1C* (Fig. 1a). These results confirm that our enriched GCPs are ABA-responsive.

**Fig. 1.**
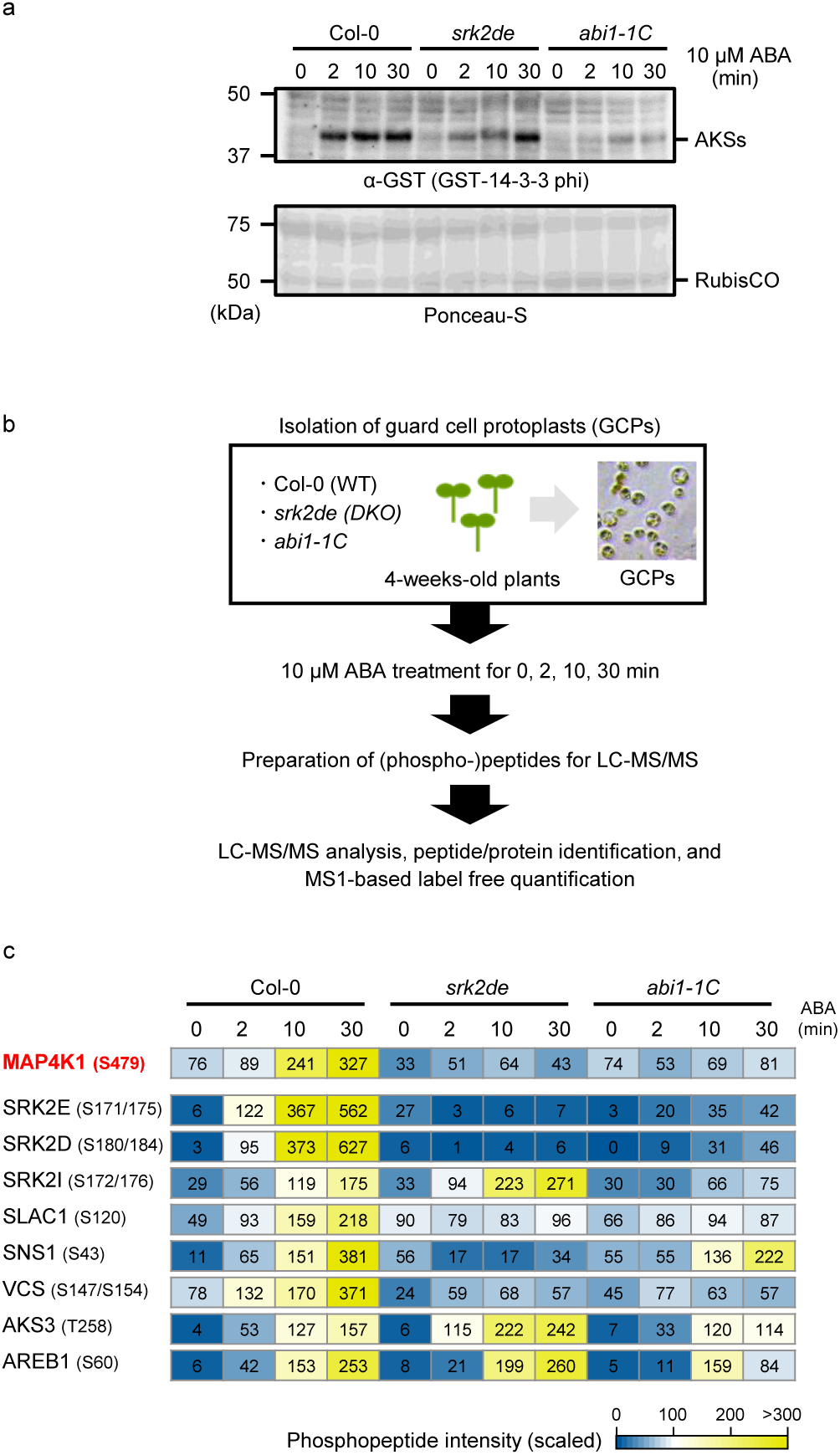
An overview of phosphoproteomic analysis using Arabidopsis guard cells. **a,** Far-western blotting analysis of 14-3-3 protein. Protein extracts of guard cell protoplasts (GCPs) were prepared from Col-0, *srk2de*, and *abi1-1C* treated with 10 µM ABA. GST-tagged 14-3-3 (GF14 phi) protein was used as a probe. Black arrows and open arrows indicate the position of AKSs and RubisCO, respectively. This experiment is repeated four times biologically independent with similar results. **b,** Workflow of the phosphoproteomic analysis for Arabidopsis GCPs. **c,** Heat map showing the intensity of phosphopeptides from known or putative SnRK2 substrates. The mean intensities were scaled to the same range for each phosphopeptide.

For phosphoproteomic analysis, samples were prepared from Col-0, *srk2de* and *abi1-1C* GCPs for LC-MS/MS analysis (Fig. 1b). From all GCP samples, we identified a total of 10,356 protein groups (Supplemental table 1) which overlapped with 94.5% of protein groups previously identified in Arabidopsis GCPs ^22^ (Fig. S1a). In our data, 1,894 phosphopeptides were significantly upregulated or downregulated after ABA treatment in Col-0 GCPs (Fig. S1b, Supplemental Table 2). Among these phosphopeptides, 1,202 were also significantly changed in *srk2de* or *abi1-1C* GCPs compared to Col-0 GCPs (Fig. S1b, Supplemental Table 2). GO analysis of proteins associated with significantly altered phosphopeptides revealed an enrichment of several GO terms, including “stomatal movement (GO:0010119)”, “chromatin remodeling (GO:0006338)”, and “cytoskeletal organization (GO:0007010)” (Fig. S1c, Supplemental table 3).

Several phosphopeptides derived from the activation loop of subclass III SnRK2s accumulated to significantly higher levels in response to ABA (Fig.1c). However, consistent with Fig. 1a, these same phosphopeptides did not increase in abundance in GCP samples from *srk2de* or *abi1-1C*. In addition to these SnRK2-derived phosphopeptides, phosphopeptides from several known SnRK2 substrate proteins, including SLAC1, AKS3/FBH4, ABF1/AREB2, SnRK2-Substrate 1 (SNS1) and VARICOSE (VCS) ^14^, differentially accumulated in ABA-treated GCPs isolated from Col-0, suggesting that our data broadly captured ABA-induced phosphorylation events in guard cells and therefore could be useful for identifying new SnRK2 substrates (Fig. 1c).

### SRK2E/OST1 directly phosphorylates MAP4K1 at Ser-479

Among the phosphopeptides identified in ABA-treated GCPs, a phosphopeptide from MAP4K1/AtMAP4Kα1 exhibited a similar trend to known SnRK2 substrates, suggesting MAP4K1 may be a SnRK2 substrate (Fig. 1c). MAP4K1 is a member of the MAP4K gene family (Fig. S2a), and MAP4K2 is approximately 80% identical to MAP4K1 ^23^. Our dataset contained six phosphorylation sites in MAP4K1 and three phosphorylation sites in MAP4K2 (Fig. 2a and S2b). Ser-479 of MAP4K1 was significantly upregulated in response to ABA in Col-0, but not in *srk2de* and *abi1-1C* (Fig. 2b). Other phosphorylation sites, such as Ser-420 or Ser-654, remained unchanged (Fig. 2b). Ser-488 of MAP4K2 is located in a similar position relative to Ser-479 of MAP4K1, but a corresponding phosphopeptide was not detected in this study (Fig. S2b).

**Fig. 2.**
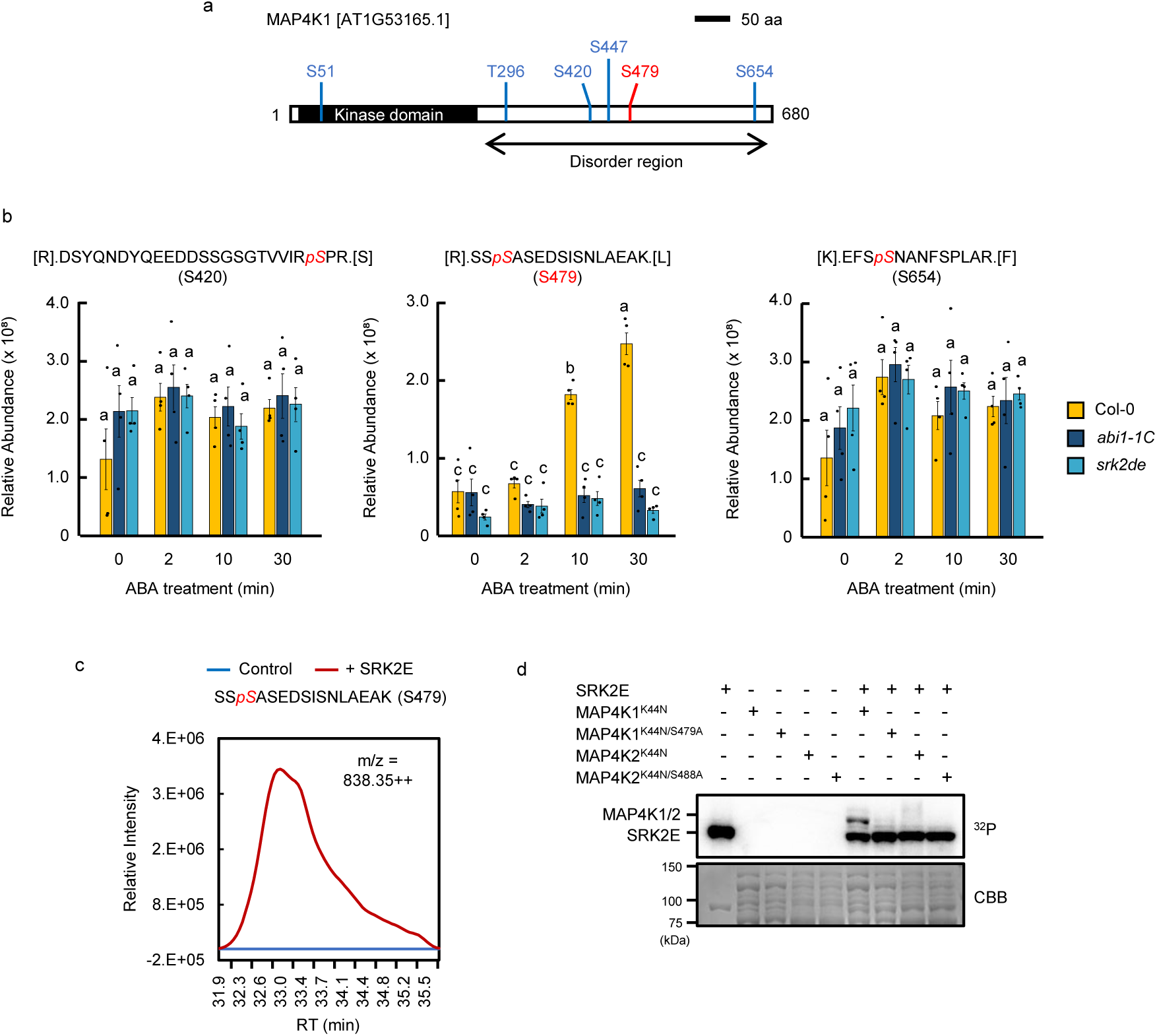
SRK2E phosphorylates MAP4K1 at Ser-479. **a,** Domain structure of Arabidopsis MAP4K1. Phosphorylation sites detected in this study are depicted as red or blue lines. **b,** Relative abundance of MAP4K1 peptides containing phosphorylated Ser-420, Ser-479, or Ser-654. Data represent means ± SE (n = 4, biologically independent replicate). Different letters indicate significant differences (Tukey’s test, P < 0.05). **c,** *In vitro* phosphorylation assay was conducted using a kinase-dead form of MBP-MAP4K1 (MAP4K1^K44N^) with or without MBP-tagged SRK2E, followed by LC-MS/MS analysis. The extracted ion chromatogram (XIC) of a MAP4K1 peptide [SS*p*SASEDSISNLAEAK] containing phosphorylated Ser-479 in the presence or absence of SRK2E. **d,** *In vitro* phosphorylation assay for SRK2E and MAP4K1/2. MBP-SRK2E was incubated with MBP-MAP4K1/2^K44N^, -MAP4K1^K44N/S479A^, or -MAP4K2^K44N/S488A^ in the presence of [γ-^32^P]ATP. Phosphorylation levels were detected by autoradiography. CBB staining showed protein loading in each lane.

To investigate whether MAP4K1 and MAP4K2 are direct substrates of SRK2E (OST1/SnRK2.6), an *in vitro* phosphorylation assay was performed by incubating *E. coli*-expressed maltose-binding protein (MBP)-tagged SRK2E together with MAP4K1^K44N^ or MAP4K2^K44N^ in a kinase reaction buffer. After incubation, the proteins were analyzed by LC-MS/MS, revealing that MBP-SRK2E phosphorylated MAP4K1 on Ser-479 (Figs. 2c). In a separate *in vitro* phosphorylation assay, MBP-MAP4K1^K44N^ was phosphorylated by MBP-SRK2E, but MAP4K1^K44N/S479A^ was not (Fig. 2d), confirming that SRK2E phosphorylates Ser-479 of MAP4K1. We also detected SRK2E phosphorylation of MAP4K2 at Ser-488 (Fig. S2c). However, the level of SRK2E phosphorylation of MBP-MAP4K2^K44N^ was lower than that of MBP-MAP4K1^K44N^ (Figs. 2d).

### IP-MS analysis of MAP4K1-interacting proteins

To identify MAP4K1-interacting proteins, we next did immunoprecipitation - mass spectrometry (IP-MS) analysis using *35S:MAP4K1-GFP* or *35S:GFP* transgenic lines. In this analysis, several SnRK2s were detected, suggesting that MAP4K1 preferentially interacts with subclass III SnRK2s (SRK2D/E/I) (Fig. 3a, Supplemental table 4). Bimolecular fluorescence complementation (BiFC) assays confirmed that SRK2E interacts with MAP4K1 and MAP4K2 in cytosol of *N. benthamiana* epidermal cells (Fig. 3b). These data are consistent with the subcellular localization of MAP4K1-GFP and SRK2E-GFP in cytosol of Arabidopsis mesophyll cell protoplasts (Fig. S3). In addition, the interaction between SRK2E and MAP4K1/2 was further supported by *in vitro* pull-down assays using MBP-MAP4K1/2 and glutathione S-transferase (GST)- tagged SRK2E (Fig. 3c).

**Fig. 3.**
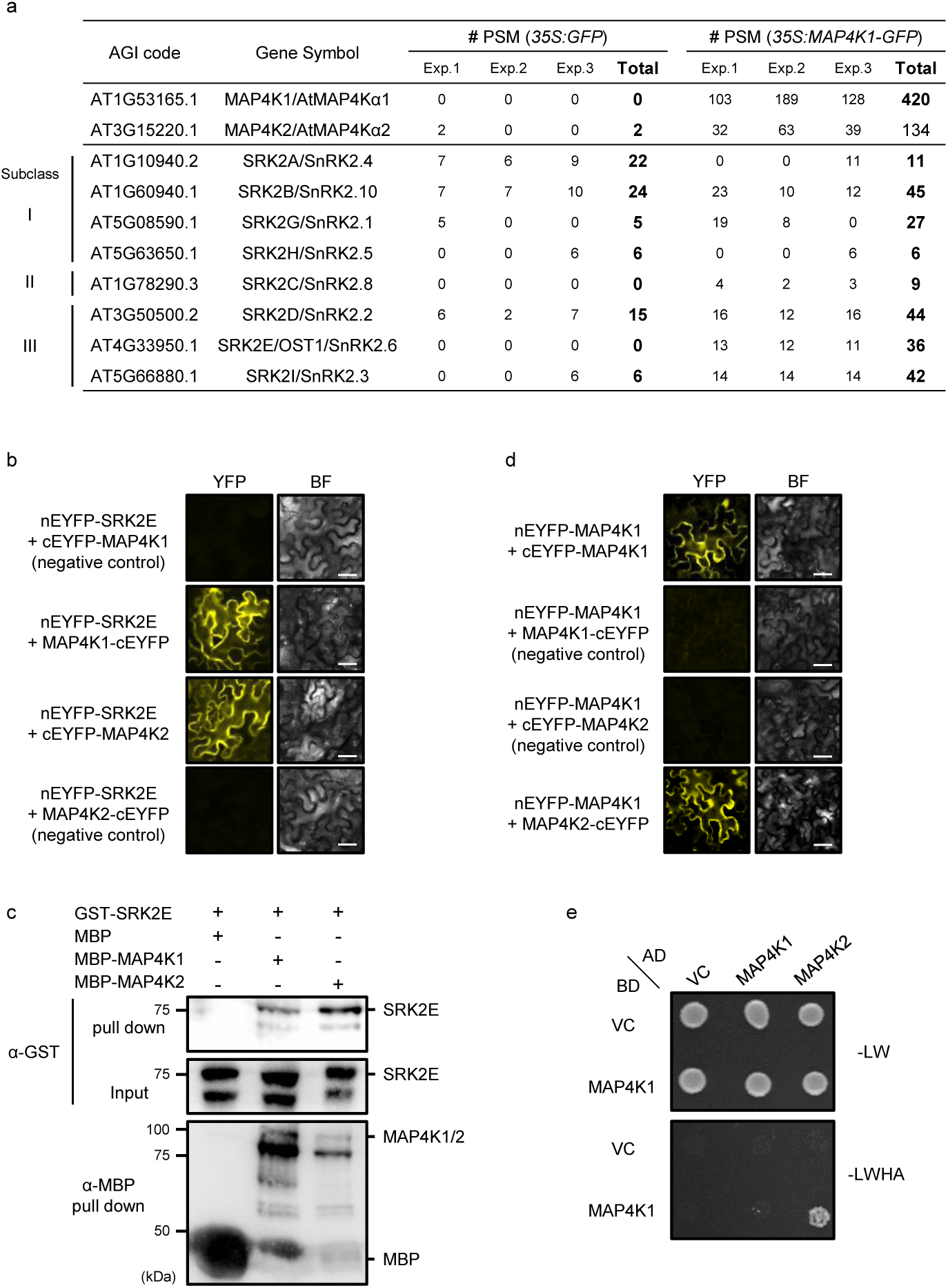
MAP4K1 interacts with MAP4K2 and subclass III SnRK2s. **a,** Immunoprecipitation (IP)-MS analysis of *35S:GFP* and *35S:MAP4K1-GFP* transgenic plants. Immunoprecipitation was performed using an anti-GFP antibody, followed by LC-MS/MS analysis. Peptide spectrum match (PSM) score indicates the number of each phosphopeptide detected in this experiment. **b,** BiFC assays were conducted for SRK2E and MAP4K1/2 in *N. benthamiana* epidermal cells. BF indicates bright field images. Scale bars indicate 50 μm. **c,** *In vitro* pull-down assay for MBP-MAP4K1/2 and GST-tagged SRK2E. GST-SRK2E and MBP-MAP4K1/2 were detected by immunoblotting using an anti-GST antibody and an anti-MBP antibody, respectively. **d,** BiFC assays between MAP4K1 and MAP4K2 in *N. benthamiana* epidermal cells. nEYFP and cEYFP represent the N- and C-terminal fragments of the enhanced yellow fluorescent protein (EYFP), respectively. BF indicates brightfield images. Scale bars indicate 50 μm. **e,** Y2H assay between MAP4K1 and MAP4K2. Yeast cells were grown on non-selective SD-Leu/ -Trp (-LW) or selective SD-Leu/ -Trp/ -His/ -Ade (- LWHA) media at 30°C for 4 days.

In addition to SRK2E-MAP4K interactions, the IP-MS analysis also detected interaction between MAP4K1 and MAP4K2 (Fig. 3a, Supplemental table 4). A BiFC assay and yeast two-hybrid (Y2H) assay confirmed that MAP4K1 and MAP4K2 directly interact each other (Fig. 3d and e). Taken together, our results reveal that MAP4K1 interacts with subclass III SnRK2s and MAP4K2 *in planta*.

### MAP4K1 and MAP4K2 redundantly function in ABA-induced stomatal closure

To gain insight into the physiological roles of MAP4K1, we tested a T-DNA insertion mutant carrying a loss-of-function allele of *MAP4K1* (*map4k1; SALK_060372*) and transgenic plants overexpressing *MAP4K1-GFP* (*35S:MAP4K1-GFP*/Col-0) for ABA-related phenotypes (Fig. S4a-c). Water loss and stomatal aperture of these plants were measured to assess stomata function. The *map4k1* mutant exhibited slightly faster water loss and a larger stomatal aperture than Col-0 (Fig. 4a-c), indicating that MAP4K1 is involved in the regulation of stomatal closure (Fig. 4a and b). On the other hand, *35S:MAP4K1-GFP*/Col-0 did not exhibit any phenotype in stomatal closure. Next, double knockout mutants of MAP4K1 and MAP4K2, *map4k1map4k2-1* and *map4k1map4k2-2,* were generated to assess their functional redundancy (Fig. S4a and b). Compared to Col-0 or single knockout mutants, the rosette leaves of *map4k1map4k2-1* and *map4k1map4k2-2* double mutants showed a more pronounced shrinkage 6 h after detachment (Fig. S4d). At 2 hours after detachment, rosette leaves from *map4k1* had a 10% increase in water loss compared to Col-0, whereas the *map4k1map4k2-1* and *map4k1map4k2-2* exhibited an approximately 20% increase in water loss (Fig. 4a). In support of this result, the aperture of stomata in *map4k1map4k2-1* and *map4k1map4k2-2* leaves was wider than that of stomata in Col-0 and *map4k1* leaves (Fig. 4b,c). These results suggest that MAP4K1 and MAP4K2 function redundantly as positive regulators of ABA-induced stomatal closure.

**Fig. 4.**
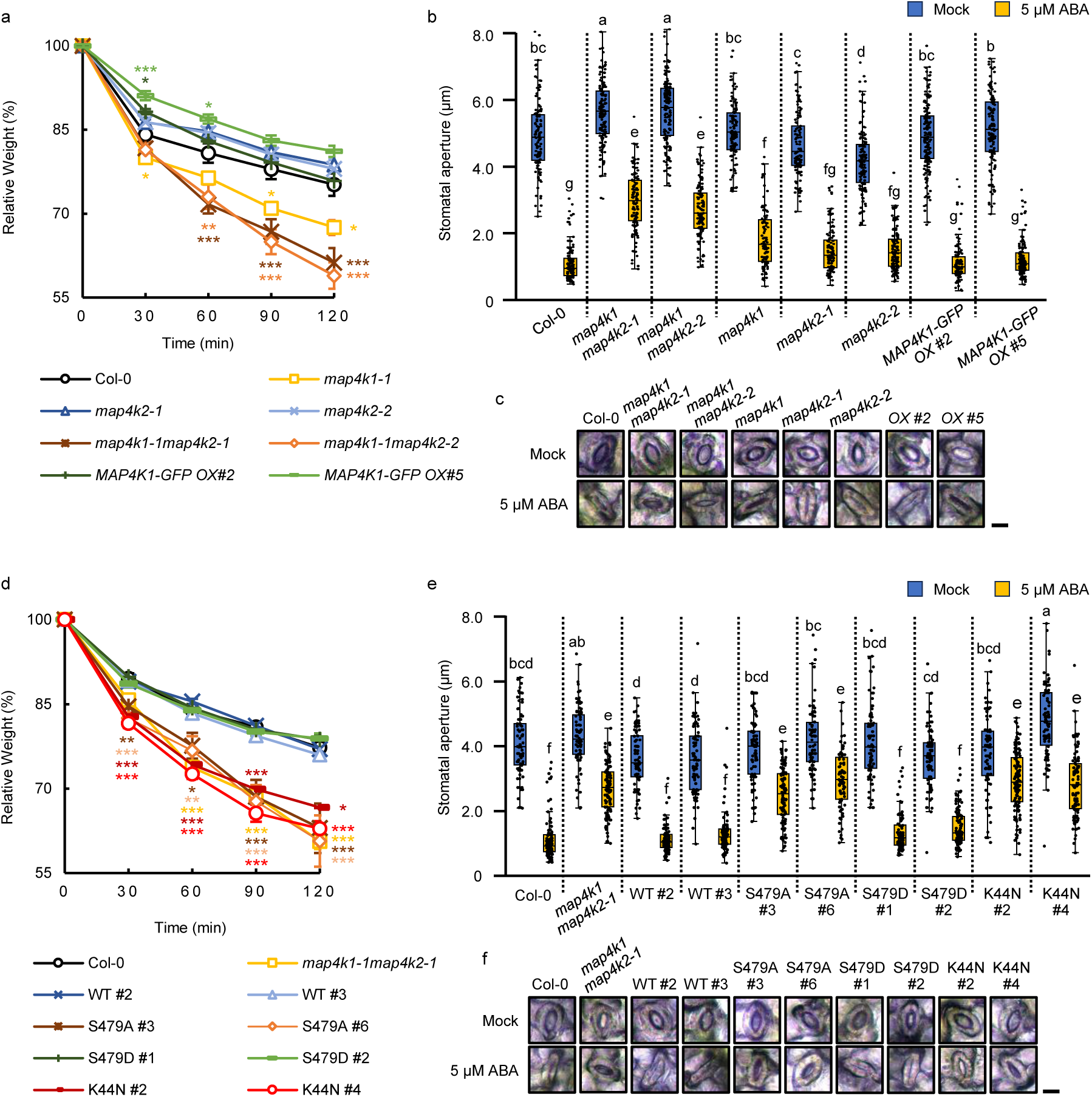
MAP4K1 and MAP4K2 positively regulate ABA-induced stomatal closure. **a,** Water loss from detached leaves of Col-0, m*ap4k1*, *map4k2-1/-2*, and *MAP4K1-GFP* overexpressing plants. Data are means ± SE (n = 3), and asterisks indicate significant differences as determined by Dunnett’s test (\**P* < 0.05, \*\**P* < 0.01, \*\*\**P* < 0.001). Each replicate consists of five individual leaves. **b,** Measurement of the stomatal aperture of Col-0, m*ap4k1*, *map4k2-1/-2*, and *MAP4K1-GFP* overexpressing plants in the presence or absence of 5 μM ABA. The data was presented as box plots; box limits represent the first and third quartiles, with the medians marked as horizontal lines. Black dots indicate raw data points from the six individual leaves of each plant. The whiskers extend up to 1.5 times the interquartile range (IQR) from the first and third quartiles, with data points beyond this range displayed as outliers. Different letters indicate significant differences (Tukey’s test, P < 0.01). **c,** Images show the representative stomata of Col-0, m*ap4k1*, *map4k2-1/-2*, and *MAP4K1-GFP* overexpressing plants. **d,** Water loss from detached leaves of Col-0, *map4k1map4k2-1*, and transgenic plants expressing FLAG-tagged MAP4K1 or its mutated forms (S479A, S479D, or K44N) in *map4k1map4k2-1* background. Each replicate consists of five individual leaves. **e,** Measurement of stomatal aperture of Col-0, *map4k1map4k2-1*, and transgenic plants expressing FLAG-tagged MAP4K1 or its mutated forms, in the presence or absence of 5 μM ABA. **f,** Images show the representative stomata of Col-0, *map4k1map4k2-1*, and transgenic plants expressing FLAG-tagged MAP4K1 or its mutated forms.

Gene expression analysis revealed that *MAP4K1* and *MAP4K2* were expressed in both GCPs and mesophyll cell protoplasts (MCPs). β-glucuronidase (GUS) reporter assays, *MAP4K1* and *MAP4K2* had similar tissue-specific expression patterns (Fig. S5a,b). In both MCPs and GCPs, the relative gene expression level of *MAP4K1* was much higher than of *MAP4K2* (Fig. S5c). *MAP4K1* was significantly induced by osmotic stress, while *MAP4K2* was not (Fig. S5d). Similar results were observed in an ABA-deficient mutant *aao3*, suggesting that *MAP4K1/2* expression is not responsive to ABA (Fig. S5d).

### Phosphorylation of MAP4K1 at Ser-479 is required for ABA-induced stomatal closure

To investigate the functional significance of phosphorylation of MAP4K1 at Ser-479, we generated mutated forms of *MAP4K1* that code for MAP4K1 with Ala and Asp substitutions at Ser-479. These S479A and S479D substitutions are expected to be non-phosphorylatable or phosphomimetic forms of MAP4K1, respectively. Additionally, we generated a kinase-dead version of *MAP4K1* that codes for MAP4K1 with Asn substituted for Lys at position 44. The altered transgenes, *MAP4K1^S479A^*, *MAP4K1^S479D^*, and *MAP4K1^K44N^*, as well as *MAP4K1^WT^*, were stably-expressed in *map4k1map4k2-1* plants. Expression of *MAP4K1^WT^* or *MAP4K1^S479D^* suppressed the enhanced water loss the phenotype of *map4k1map4k2-1* (Fig. 4d). In contrast, expression of *MAP4K1^S479A^* or *MAP4K1^K44N^*did not alter the enhanced water loss of *map4k1map4k2-1* (Fig. 4d). Consistent with these results, expression of *MAP4K1^WT^* or *MAP4K1^S479D^*rescued ABA-induced stomatal closure of *map4k1map4k2-1*, but *MAP4K1^S479A^* and *MAP4K1^K44N^*did not (Fig. 4e,f). These results suggest that the phosphorylation of MAP4K1 at Ser-479 and MAP4K1 kinase activity play a significant role in ABA-induced stomatal closure.

### MAP4K1/2 is involved in [Ca^2+^]_cyt_ elevation to regulate ABA-induced stomatal closure

SRK2E regulates ABA-induced stomatal closure via ROS production, [Ca^2+^]_cyt_ elevation, and SLAC1 phosphorylation ^4,5^. To investigate if MAP4K1/2 are required for these ABA-induced signaling events, we first applied H_2_O_2_ and Ca^2+^ to the epidermis of Arabidopsis leaves to mimic ROS production and [Ca^2+^]_cyt_ elevation, respectively. We then measured stomatal aperture in the treated leaves. Consistent with previous studies ^7,12,24^, ABA, H_2_O_2_ and Ca^2+^ induced stomatal closure in Col-0 leaves (Fig. 5a). In *map4k1map4k2*, stomatal closure in response to ABA and H_2_O_2_ was completely impaired, but was normally induced by Ca^2+^ (Fig. 5a). This result suggested that MAP4K1/2 functions upstream of [Ca^2+^]_cyt_ elevation. ABA-responsive ROS production in guard cells was observed at a similar level in Col-0 and *map4k1map4k2-1* but reduced in *srk2e/ost1-3* (Fig. 5b,c). The reduced response of *srk2e/ost1-3* was specific to ABA treatment, as similar levels of light-induced ROS production occurred in Col-0, *map4k1map4k2-1* and *srk2e/ost1-3* (Fig. 5b,c). These data suggest that MAP4K1/2 may not be involved in light- or ABA-responsive ROS production. Next, the relationship between MAP4K1/2 and SLAC1 was examined. Although MAP4K1/2 did not directly phosphorylate the N-terminus and C-terminus of SLAC1 *in vitro* (Fig. S6a), phosphorylation of SLAC1 at Ser-86, which is essential for SLAC1 activation ^25,26^, was significantly decreased in GCPs of *map4k1map4k2-1* (Fig. S6b,c). These results suggest that MAP4K1/2 affects SLAC1 phosphorylation in response to ABA but not through direct phosphorylation.

**Fig. 5.**
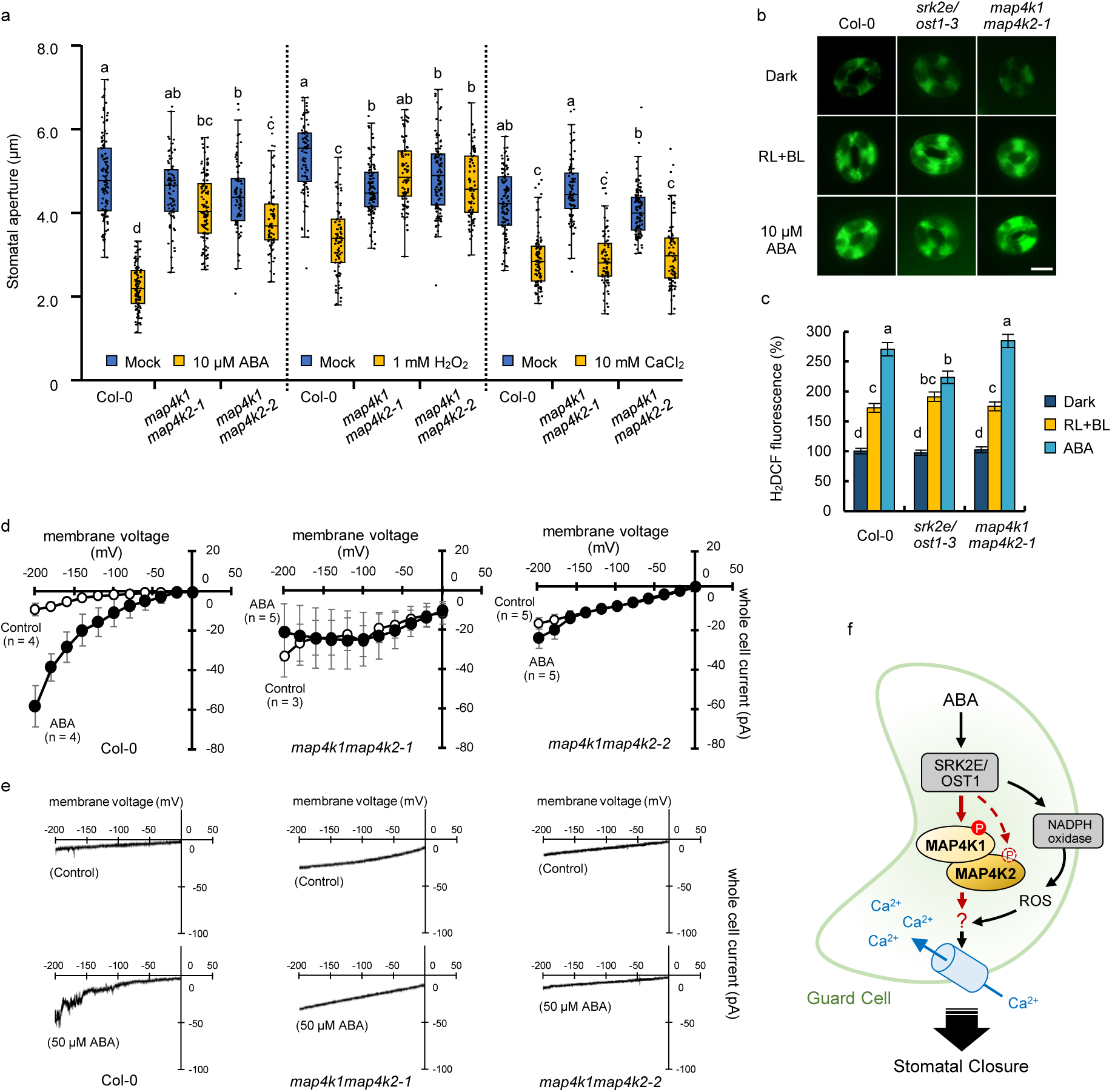
MAP4K1 and MAP4K2 are involved in the activation of plasma membrane Ca^2+^-permeable (*I_Ca_*) channels in ABA-induced stomatal closure. **a,** Measurement of the stomatal aperture of Col-0, *map4k1map4k2-1* and *map4k1map4k2-2*. Stomatal aperture was presented as box plots; the box limits represent the first and third quartiles, with the medians marked as horizontal lines. Black dots indicate the raw data points from the six individual leaves of each plant. The whiskers extend up to 1.5 times the interquartile range (IQR) of the first and third quartiles, with data points beyond this range displayed as outliers. Different letters indicate significant differences (Tukey’s test, P < 0.01). **b,c,** ROS accumulation in guard cells were indicated by the fluorescent dye H_2_DCFDA (b), and the relative ROS levels were quantified using ImageJ software (c). Epidermal strips were incubated in the dark or under red light (RL) and blue light (BL), and then loaded with H_2_DCFDA. 10 μM ABA was added and reactions for another 30 min. The scale bar indicates 10 μm. Data represent means ± SE (n = 75, pooled from triplicate experiments). Different letters indicate significant differences (Tukey’s test, P < 0.05). **d,** Average current-voltage curves for ABA-activated *I_Ca_* channels in GCPs isolated from Col-0, *map4k1map4k2-1*, and *map4k1map4k2-2*. The data represent mean ± SE. Open and closed circles indicate mock and 50 µM ABA treatment, respectively. ABA was added approximately 16 min after the start of whole-cell recording. Voltage was ramped from 0 to −198 mV at a ramp rate of 100 mV s^−1^. **e,** Representative whole-cell Ca^2+^ current recordings from Col-0, *map4k1map4k2-1* and *map4k1map4k2-2* GCPs. **f,** A model of the SRK2E-MAP4K1/2 pathway during ABA-induced stomatal closure. ABA-activated SRK2E directly phosphorylates Ser-479 of MAP4K1 to regulates ABA-induced stomatal closure through activation of *I_Ca_* channels. MAP4K1/2-regulated *I_Ca_* channels may share or partially overlap with ROS-regulated.

To measure ABA-dependent activation of plasma membrane Ca^2+^-permeable (*I_Ca_*) channels, a whole-cell patch clamp recording was performed using GCPs. In agreement with previous reports ^11,27,28^, externally applied ABA activated *I_Ca_* channel currents in Col-0 GCPs (Fig. 5d,e). However, ABA activation of *I_Ca_* channels was not observed in *map4k1map4k2-1* or *map4k1map4k2-2* GCPs (Fig. 4d,e). These results further support the involvement of MAP4K1/2 in ABA-responsive elevation of [Ca^2+^]_cyt_ in guard cells.

Our data support two possible models for how MAPK4K1/2 contributes to ABA-induced elevation of [Ca^2+^]_cyt_. First, MAP4K1/2 may function as an intermediary between SnRK2 and *I_Ca_* channels. Alternatively, MAP4K1/2 may regulate SnRK2 activity, which in turn influences [Ca^2+^]_cyt_. To test the latter model, an *in vitro* phosphorylation assay was conducted using recombinant MAP4K1/2 and a kinase-dead form SRK2E^K50N^. In this assay, MAP4K1/2 did not phosphorylate SRK2E^K50N^ (Fig. S7a). AtARK1/Raf4, an upstream protein kinase of SnRK2^29–31^, was used as a positive control. To examine the possibility that MAP4K1/2 indirectly affects SnRK2 activity, *in-gel* kinase assays and 14-3-3 far-western blot analyses using both seedlings and GCPs were performed. The levels of SnRK2 activity was similar between Col-0 and *map4k1map4k2-1* (Fig. S7b-e). Taken together, our results indicate that MAP4K1/2 functions downstream of SnRK2s and regulates [Ca^2+^]_cyt_ elevation via *I_Ca_* channels.

## Discussion

SRK2E (OST1/SnRK2.6), a member of subclass III SnRK2s, plays a critical role in ABA-induced stomatal closure, a process that is essential for plants to tolerate drought ^4,5^. To identify SnRK2 substrates, we did a phosphoproteomic analysis using Arabidopsis guard cell protoplasts (GCPs) (Fig. 1b). The resulting dataset contains a total of 10,356 protein groups from Col-0, *srk2de*, and *abi1-1C* (Supplemental table 1). This scale is comparable to a previous study on Arabidopsis guard cells ^22^ (Fig. S1a). Furthermore, GO analysis confirmed that our data retained protein profiles that are known to occur in Arabidopsis guard cells (Supplemental table 3, Fig. S1c). These results suggest that our phosphoproteomic data is suitable for screening SnRK2 substrates in guard cells. In this regard, we successfully identified MAP4K1 as a substrate of SnRK2 through phosphoproteomics-based screening (Fig. 1c). In general, to identify a protein (X) as a bona fide substrate of a protein kinase (Y), certain criteria should be met. For instance, X should be phosphorylated under the regulation of Y *in vivo*, and Y should directly phosphorylate X *in vivo* or *in vitro*. In the case of MAP4K1, our phosphoproteomic analysis revealed that Ser-479 of MAP4K1 is phosphorylated in Arabidopsis guard cells in response to ABA (Fig. 2b), and several lines of experiments showed that SRK2E can phosphorylate MAP4K1 at Ser-479 *in vivo* or *in vitro* (Fig. 2 and Fig 3). Thus, MAP4K1 meets the criteria to be considered a substrate phosphorylated by SnRK2.

Our genetic analyses revealed that MAP4K1 positively regulates ABA-induced stomatal closure in Arabidopsis (Fig. 4a-c), and that phosphorylation of Ser-479 is required for MAP4K1 function (Fig. 2d-f). These findings suggests that a phospho-signaling pathway involving SnRK2 and MAP4K1 could be operational in response to ABA. In addition, MAP4K2 is the closest relative of MAP4K1 and is functionally redundant with MAP4K1 in regulating ABA-induced stomatal closure (Fig. 2). Both of MAP4K1 and MAP4K2 directly interacted with SRK2E (Fig. 3a and b), suggesting that this shared interaction might be the cause of the redundancy. Although Ser-488 of MAP4K2 can be phosphorylated by SnRK2 *in vitro* (Fig S2c), it is still unclear whether MAP4K2 is phosphorylated *in vivo* in response to ABA (Fig. S2b).

IP-MS and biochemical experiments revealed that MAP4K1 interacts with MAP4K2 (Fig. 3a,d,e. In Arabidopsis, MAP4K4/TARGET OF TEMPERATURE3 (TOT3) physically interacts with MAP4K5/TOT3-INTERACTING PROTEIN5 (TOI5) and MAP4K6/TOI4, and TOT3 functions redundantly with TOI5/TOI4 in hypocotyl/petiole elongation during thermomorphogenesis ^32^. Furthermore, BLUE LIGHT SIGNALING1 (BLUS1)/MAP4K10 is directly phosphorylated by PHOT1/2 in a blue light-dependent manner, releasing BLUS1 from autoinhibition^33–35^. In our experiments the kinase activity of MAP4K1 or MAP4K2 was not affected by Asp substitution of Ser-479 or Ser-488, respectively (Fig. S8). Though, in some instances Ser to Asp substitution does not adequately mimic a phosphorylated Ser. Therefore, further studies will be required to understand the role of MAP4K1/2 phosphorylation in MAP4K1/2 activation and ABA signaling.

Our data demonstrate that MAP4K1/MAP4K2 are genetically required for ABA-induced increases in [Ca^2+^]_cyt_ and stomatal closure. The driving force behind stomatal closure is ion efflux, and one of the major regulatory pathways involves SnRK2 directly phosphorylating S-type anion channels including SLAC1 ^4,5^. Ca^2+^-dependent pathways are also involved in the regulation of SLAC1 ^28^. In this regard, ABA-induced ROS triggers an increase in [Ca^2+^]_cyt_ ^6,7^, which in turn activates CDPKs or CBLs-CIPKs to phosphorylate S-type anion channels ^27,36^. By directly measuring the activity of *I_Ca_* channels, we discovered that MAP4K1/2 is required for ABA-induced increases in [Ca^2+^]_cyt_ (Fig. 5c). ABA-responsive activation of *I_Ca_* was largely impaired in *map4k1map4k2* (Fig. 5d and e), whereas ABA-dependent ROS production occurred at normal levels in *map4k1map4k2* (Fig. 5b,c). These data imply that MAP4K1/2 is not involved in ROS production but rather functions upstream of [Ca^2+^]_cyt_ elevation. Notably, the phosphorylation level of SLAC1 at Ser-86, which is critical for anion channel activity^25,26^, was significantly decreased in *map4k1map4k2-1* (Fig. S6c). Taken together, our study proposes that the signaling module of SnRK2-MAP4K1/2 functions to regulate ABA-induced stomatal closure via a Ca^2+^-dependent signaling pathway (Fig. 5f). However, the mechanism(s) by which MAP4K1/2 regulates [Ca^2+^]_cyt_ and *I_Ca_* is currently unknown. Recently, some (pseudo-)protein kinases, such as GUARD CELL HYDROGEN PEROXIDE-RESISTANT1 (GHR1) and CANNOT RESPOND TO DMBQ1/H_2_O_2_-induced Ca^2+^ increases 1 (CARD1/HPCA1) ^37–40^, have been proposed to be involved in the ROS-dependent regulation of [Ca^2+^]_cyt_ in guard cells. In addition, the ABA-activated *I_Ca_* channels CYCLIC NUCLEOTIDE-GATED CHANNEL (CNGC) 5/6/9/12 function in ROS-independent [Ca^2+^]_cyt_ elevation in guard cells ^41^, and Ser-27 at the N-terminus of CNGC6 is phosphorylated by SRK2E^42^. Therefore, it is likely that *I_Ca_* channels in guard cells are regulated through both ROS-dependent and -independent pathways. Further studies will be required to clarify the relationship between MAP4K1/2 and the regulatory pathways of *I_Ca_* channels.

Notably, MAP4K1/2 likely functions not only in guard cells but also in other plant tissues. For example, *MAP4K1/2* is also expressed in leaves and roots (Fig. S5a and b), and *map4k1map4k2* seedlings showed ABA insensitivity in cotyledon greening rate and stress-responsive gene expression (Fig. S9a and b), indicating that MAP4K1/2 are functional at this early stage of development. Furthermore, MAP4K1 was phosphorylated in response to ABA in Arabidopsis seedlings (Fig. S9c), suggested that MAP4K1 may contribute to ABA signaling in tissues other than guard cells. This point might make difficult to find MAP4K1/2 in previous studies on guard cell-specific factors. In homologs of MAP4K1 in other plant species, Ser residues corresponding to Ser-479 of MAP4K1 are well conserved among dicots, with the exception of legumes (Fig. S10) ^43^. This suggests that SnRK2-dependent phosphorylation of MAP4K1/2 may occur in a diversity of dicotyledonous crop plants. Further elucidation of MAP4K1/2 functions and signaling mechanisms will provide new insights into ABA responses in plants, and may provide new avenues for future targeted engineering of drought tolerance or water use efficiency of crop species.

## Methods

### Plant materials and growth conditions

Arabidopsis *thaliana* ecotype Columbia (Col-0) was used as wild-type plants. T-DNA insertion mutant lines, *map4k1* (SALK_060372), *map4k2-1* (SAIL_1255_G11), and *map4k2-2* (*SALK_115951*), were obtained from the Arabidopsis Biological Resource Center (ABRC). The double mutants, *map4k1map4k2-1* and *map4k1map4k2-2* were generated by crossing *map4k1* and map4k2-*1* or *map4k2-2*. ABA response mutants, *srk2e/ost1-3, srk2de*, and *abi1-1C*, were obtained as previously described ^19^. Plant growth conditions were described in a previous study ^44^. To test responses to ABA or osmotic stress, seeds were sown on GM medium supplemented with or without indicated concentrations of ABA (Sigma-Aldrich) or mannitol (Wako), and cotyledon greening rates were recorded for two weeks after stratification.

### Isolation of Arabidopsis guard cell protoplasts (GCPs) and ABA treatment

Arabidopsis guard cell protoplasts (GCPs) were enzymatically isolated from rosette leaves of 4- to 5-weeks-old plants, following procedures outlined in previous studies ^17,18^. GCPs were adapted to the dark for 3 h at 4°C, and then they were incubated in H^+^-pumping buffer [0.125 mM MES-NaOH (pH 6.0), 1 mM CaCl_2_, 0.4 M mannitol, and 10 mM KCl] under white light [90 µmol m^-2^ s^-1^] for 1 h at 24°C. Samples were incubated with 10 µM ABA for the indicated periods. The reaction was then stopped by adding trichloroacetic acid (TCA) for far-western blotting, or by using liquid nitrogen for phosphoproteomic analysis, as described below.

### GST-14-3-3 far-western blotting

Far-western blotting analysis was performed using GST-tagged 14-3-3phi (GF14phi) protein as a probe according to a previous study with slight modifications ^20,21^. In brief, precipitated GCP proteins were resuspended in sample loading buffer [62.5 mM Tris-HCl (pH 6.8), 2 % SDS, 5 % sucrose, 2 ppm bromophenol blue, 100 mM dithiothreitol (DTT)] and separated on 10% SDS-PAGE. After electrophoresis, the proteins were transferred to the polyvinylidene fluoride (PVDF) membrane (0.45 μm, Millipore) and then incubated with GST-14-3-3, which were detected using anti-GST HRP conjugate antibody (RPN1236, Cytiva, MA).

### Vector constructions

The full-length cDNAs of *MAP4K1* and *MAP4K2* were cloned into pENTR1A vector (Thermo Fisher Scientific) and confirmed by sequencing. Amino acid substitutions were carried out using site-directed mutagenesis as previously described ^45^. The cDNAs were then transferred into destination vectors, such as pGreen0029-GFP ^19^, R4pGWB 501 ^46^, pSITE-cEYFP-C1 (CD3-1649) or pSITE-cEYFP-N1 (CD3-1651), using Gateway LR Clonase II (Thermo Fisher Scientific). 1300 or 1928 bp upstream sequences of *MAP4K1* or *MAP4K2*, respectively, were amplified with specific primers with Gateway attB4/attB1r adaptor sequences and cloned into pDONR P4-P1R vector (Thermo Fisher Scientific) using Gateway BP Clonase II (Thermo Fisher Scientific) to generate pDONR P4-P1R *MAP4K1p* and pDONR P4-P1R *MAP4K2p*. pDONR P4-P1R *MAP4K1p* or pDONR P4-P1R *MAP4K2p*, along with pENTR-GUS (Thermo Fisher Scientific), were combined with R4pGWB 501 ^46^ using Gateway LR Clonase II to generate R4pGWB 501 *MAP4K1p:GUS* and R4pGWB 501 *MAP4K2p:GUS*.

### Transgenic plants

The pGreen0029-GFP construct harboring *MAP4K1* was utilized to overexpress MAP4K1-GFP, was prepared as described above. R4pGWB 501 constructs harboring *MAP4K1p:GUS, MAP4K2p:GUS*, and *MAP4K1p:MAP4K1^WT^/MAP4K1^S479A^/MAP4K1^S479D^/MAP4K1^K44N^-3xFLAG* were prepared as well. Each construct was transformed into Col-0 or *map4k1map4k2-1* with *A. tumefaciens* strain GV3101 or GV3101 (with pSOUP). Transgenic plants were selected on GM medium containing 200 μg/mL Claforan with 50 μg/mL kanamycin or 25 μg/mL hygromycin. The expression of GFP-tagged or FLAG-tagged proteins were confirmed by immunoblotting using anti-GFP pAb (MBL, code No. 598) or anti-FLAG (DYKDDDDK) (Wako, 014-22383, lot SAR0168) as the primary antibodies, and using horse anti-rabbit IgG antibody or horse anti-mouse IgG antibody (Vector Laboratories, CA) as the secondary antibodies, respectively.

### Transient expression and subcellular localization analysis

Preparation of Arabidopsis mesophyll cell protoplasts (MCPs) and PEG-calcium transfection were performed according to previous studies ^47^. For transient expression, pGreen0029-GFP constructs harboring *SRK2E* ^19^ or *MAP4K1* were prepared from *E. coli* (Takara, DH5α) using QIAGEN® Plasmid Midi Kit (QIAGEN) following the manufacturer’s protocol. For subcellular localization analysis, 10 µg plasmid DNA was transfected into 5 x 10^4^ protoplasts, and GFP fluorescence was observed using a BX53 fluorescence microscope (Olympus).

### Phylogenetic analysis and prediction of kinase domains

Arabidopsis MAP4K1 orthologs were determined using BLASTP search implemented in Phytozome v13 ^48^ (https://phytozome-next.jgi.doe.gov/) against the specified organisms. Multiple alignments for the full-length protein sequence were generated by MUSCLE algorithm ^49^ implemented in MEGA-X ^50^. The phylogenetic trees were built with the Neighbor-Joining method ^51^ based on the multiple alignments. All positions containing gaps were eliminated, and the phylogenies were evaluated using bootstrap with 1000 replicates ^52^. Prediction of kinase domains for each protein was performed using PROSITE ^53^ (https://prosite.expasy.org/).

### Histochemical GUS staining

Arabidopsis seedlings of *MAP4K1p:GUS* or *MAP4K2p:GUS* transgenic plants were pretreated with 90% cold acetone, then incubated in GUS staining solution, containing 1 mM 5-bromo-4-chloro-3-indolyl-β-D-glucuronic acid (X-Gluc), 0.5 mM K_3_[Fe(CN)_6_], 0.5 mM K_4_[Fe(CN)_6_], 0.1 % Triton X-100, 50 mM sodium phosphate (pH 7.2), in the dark at 37°C overnight. The samples were washed and bleached with 70% EtOH. Images of the stained samples were captured using a microscope or an image scanner.

### Water loss analysis

Arabidopsis 1-week-old plants were grown on GM agar plates under long-day conditions (16 h light/8 h dark) at 22°C were transferred to soil. The plants were then grown under the same conditions for an additional three weeks. The detached rosette leaves from approximately 4-week-old plants were placed on weighing dishes and left on the laboratory bench. Fresh weights were measured as previously described ^44^. Statistical analysis was conducted using Dunnett’s test with the R package.

### Measurement of stomatal aperture

Stomatal aperture in the abaxial epidermis was measured following previous studies with slight modifications ^54^. In brief, epidermal strips were peeled from the rosette leaves of 4 to 5-week-old Arabidopsis seedlings grown on soil. The strips were then incubated in stomatal opening buffer [10 mM MES-KOH (pH 6.2), 5 mM KCl, and 0.1 mM CaCl_2_] under white light for 3 h to fully expand the stomatal pore. Then the strips were transferred to stomatal opening buffer containing the indicated concentrations of ABA for 2 h. Stomatal apertures were photographed using a microscope and measured with ImageJ software. Statistical analysis was performed using Tuekey’s HSD test with the R package.

### RNA extraction and RT-qPCR analysis

Total RNA was extracted from 2-week-old seedlings, mesophyll cell protoplasts (MCPs) or guard cell protoplasts (GCPs) as described previously ^44^, and 500 ng of total RNA was used for reverse transcription using ReverTra Ace qPCR RT Master Mix with gDNA Remover (TOYOBO). RT-qPCR analysis was performed using LightCycler 480 SYBR Green I Master (Roche Life Science) with Light Cycler 96 (Roche Life Science). Each transcript was normalized by *GAPDH* and analyzed with three biological replicates. The gene-specific primers used for RT-qPCR are listed in Supplementary Table 5.

### Preparation of recombinant proteins

pMAL-c5X constructs harboring *SRK2E^K50N^* and *AtARK1/Raf4* cDNAs were prepared previously^29^. *MAP4K1, MAP4K2*, and *SLAC1* cDNAs (N-terminal or C-terminal) were cloned in-frame into pMAL-c5X vector (New England Biolabs) using In-Fusion HD Cloning Kit (Takara Bio). Amino acid substitutions were introduced into pMAL-c5X MAP4K1 or MAP4K2 by site-directed mutagenesis to produce the kinase-dead form of *MAP4K1^K44N^* or *MAP4K2^K44N^* and the non-phosphorylatable form of *MAP4K1^K44N/S479A^ or MAP4K2^K44N/S488A^*. The recombinant proteins were expressed and affinity purified from *E. coli* strain BL21 (DE3) using Amylose Resin (New England Biolabs), as previously described ^44^.

### *In vitro* phosphorylation assays

*In vitro* kinase assay was performed as described previously ^44^. For LC-MS/MS analysis, proteins that underwent *in vitro* reactions were digested in 50 mM ammonium bicarbonate (NH_4_HCO_3_) with 0.2 µg trypsin (Promega) at 37°C overnight. Tryptic digestion was stopped by adding trifluoroacetic acid (TFA) to a final concentration of 1 %. Peptides were desalted using an in-house C18 Stage-tip ^55^ made with a C18 Empore disk (CDS). Desalted peptides were dried using a centrifugal concentrator (CC-105, TOMY), and stored at −20°C until LC-MS/MS analysis.

### *In-gel* kinase assays

*In-gel* kinase assay was performed as previously described with slight modifications ^44^. Protein extraction buffer for 2-week-old Arabidopsis seedlings or isolated GCPs was modified as described below: 20 mM HEPES-KOH (pH 7.5), 0.5% TritonX-100, 100 mM NaCl, 5 mM MgCl_2_, 0.1 mM EDTA, 1 mM Na_3_VO_4_, 25 mM NaF, 50 mM β-glycerophosphate, 20% (v/v) glycerol, and 1% (v/v) protease inhibitor cocktail (Sigma-Aldrich).

### Sample preparation for IP-MS analysis

Crude proteins were extracted from the finely ground powder of 2-week-old seedlings of *35S:GFP* and *35S:MAP4K1-GFP* transgenic plants using a protein extraction buffer, containing 20 mM HEPES-KOH (pH 7.5), 100 mM NaCl, 0.1 mM EDTA, 5 mM MgCl_2_, 20% glycerol, 0.5% TritonX-100, 1 mM Na_3_VO_4_, 25 mM NaF, and 1% Protease Inhibitor Cocktail (Sigma-Aldrich). The mixture was then centrifuged to remove cellular debris. The supernatant was incubated with GFP-selector (NanoTag Biotechnologies) for 1.5 h at 4°C. After centrifugation, GFP-selector beads were washed four times with the protein extraction buffer and once with 50 mM NH_4_HCO_3_. The reduction and alkylation mixture (10 mM tris(2-carboxyethyl)phosphine (TCEP), 40 mM chloroacetamide (CAA), 50 mM NH_4_HCO_3_) was added to the washed beads, followed by incubation at 24°C for 30 min in the dark. Then 0.2 µg Trypsin (Promega) was added to the beads, followed by oevernight incubation at 37°C. Tryptic digestion was stopped by adding trifluoroacetic acid (TFA) to final concentration of 1 %. Peptides were desalted using an in-house C18 Stage-tip ^55^ made with a C18 Empore disk (CDS). Desalted peptides were dried using a centrifugal concentrator (CC-105, TOMY) and stored at −20°C until LC-MS/MS analysis.

### Bimolecular fluorescence complementation (BiFC) assay

BiFC assay was performed as previously described ^44^. In brief, *A. tumefaciens* strain GV3101 (p19) expressing pSITE-nEYFP-N1:*SRK2E*, pSITE-cEYFP-N1:*MAP4K1/MAP4K2*, or pSITE-cEYFP-C1:*MAP4K1/MAP4K2* were mixed as indicated pairs and infiltrated into *N. benthamiana* leaves. After a few days, complementary fluorescence of YFP was observed in epidermal cells using a BX53 fluorescence microscope (Olympus, Japan).

### *In vitro* pull-down assay

*In vitro* pull-down assay was performed as previously described ^56^. In brief, *E. coli* cells expressing MBP, MBP-MAP4K1, MBP-MAP4K2, or GST-SRK2E were homogenized with lysis buffer [50 mM Tris-HCl (pH 7.4), 150 mM NaCl, 1% NP40, 0.25% sodium deoxycholate (SDC)]. Lysates containing GST-SRK2E or MBP-tagged proteins were mixed, and a portion of the mixture was set aside as input fraction. Amylose resin beads (New England Biolabs, MA) beads were added to the mixture and incubated for 3 h at 4°C. The beads were washed four times with lysis buffer. Proteins were then eluted from the beads using lysis buffer containing 10 mM maltose. Input and eluted fractions were detected by western blotting.

### Sample preparation for phosphoproteomic/proteomic analysis

Protein extraction and digestion were performed according to a previous study with some modifications ^57^. In brief, total protein lysates were extracted from isolated GCPs in protein extraction buffer [100 mM Tris-HCl (pH 9.0), 6 M guanidine hydrochloride (Gdn-HCl)], followed by heating the lysates for 5 min at 95°C. Place the lysate samples on ice, sonicate the samples using a micro ultrasonic homogenizer (Q125, QSONICA, USA), and centrifuge at 17,400 x *g* for 20 min at 4°C. The supernatant was then precipitated by the methanol-chloroform method, and the protein pellets were resuspended in a digestion buffer containing 100 mM Tris-HCl (pH 9.0), 12 mM SLS, and 12 mM SDC. 200 μg of protein per sample was incubated in a reduction and alkylation mixture [10 mM TCEP, 40 mM CAA, 50 mM ammonium bicarbonate] for 30 min at 24°C in the dark. After a five-fold dilution with 50 mM ammonium bicarbonate, proteins were digested overnight at 37°C with 2 µg trypsin (Promega). Phase transfer surfactants (such as SLS, SDC) were removed according to a previous study ^58^. After removal, 10 % of the total volume of peptide samples was set aside for global proteomic analysis, and phosphopeptides were enriched using hydroxy acid-modified metal oxide chromatography (HAMMOC) as described in previous studies ^59,60^. Digested peptides or enriched phosphopeptides were desalted using an in-house Stage-tip made with SDB Empore disks (CDS). After desalting, the phosphopeptides were dried and stored at −20°C until LC-MS/MS analysis.

### LC-MS/MS analysis and raw data processing

Prepared peptide samples were analyzed using a nano LC system, Easy-nLC 1200 (Thermo Scientific), connected in line to a quadrupole-Orbitrap mass spectrometer, Orbitrap Exploris 480 (Thermo Scientific), equipped with an aerodynamic high-field asymmetric waveform ion mobility spectrometry (FAIMS) device, FAIMS Pro (Thermo Scientific).

Dried peptides were resuspended in a solution of 2% (v/v) acetonitrile (ACN) with 0.1% (v/v) formic acid (FA) and then loaded directly into a C18 nano HPLC capillary column (NTCC-360/75-3, 75 µm ID × 15 cm L, Nikkyo Technos). Peptides were eluted at 300 nL/min, at 60°C with nonlinear gradient program for 140 min for phosphoproteomic analysis. IP-MS and *in vitro* kinase samples were eluted for 105 min. The mobile phase buffer consisted of 0.1% FA (Buffer A) with an elution buffer of 0.1% FA in 80% ACN (Buffer B). The 140 min program was executed under the following conditions: *0-5 min, B 6% held; 5-79 min, B 6-23%; 79-107 min, B 23-35%; 107-125 min, B 35-50%; 125-130 min, B 50-90%; 130-140 min, B 90% held*. The 105 min program was executed as follows: *0-54 min, B 6-23%; 54-75 min, B 23-35%; 75-90 min, B 35-50%; 90-95 min, B 50-65%; 95-105 min, B 65% held*. Eluted peptides were ionized at a source voltage of 2.2 kV and detected using data-dependent acquisition (DDA) in positive ion mode. MS1 spectra were collected in the range of 375-1500 *m/z*. For phosphoproteomic analysis, the resolving power was set to 60,000. For IP-MS and *in vitro* kinase assay, the resolving power was set to 120,000. For all analyses, MS2 spectra were collected in the range above 120 *m/z* at a resolving power of 30,000. In the FAIMS compensation voltage (CV), two conditions were used (−40/-60 CV or −50/-70 CV) and the resolution was set to “Standard” (inner temperature (IT) 100°C / outer temperature (OT) 100°C) in both conditions.

For phosphoproteomic or proteomic analysis and *in vitro* kinase assay, peptide/ protein identification and MS1-based label-free quantification (LFQ) were performed using Proteome Discoverer 2.5 (PD2.5) (Thermo Scientific). For IP-MS analysis, peptide/ protein identification was carried out using PD3.1 (Thermo Scientific). MS/MS spectra were searched using SEQUEST HT (PD2.5) or CHIMERYS (PD3.1) algorithms against the Arabidopsis protein database (Araport11_genes.201606.pep.fasta). In both of search engines, the following search parameters were used as follows: *[Digestion Enzyme; trypsin], [Maximum Missed Cleaves; 2], [Peptide Length; 6-144], [Precursor Mass Tolerance; 10 ppm], [Fragment Mass Tolerance; 0.02 Da], [Static Modification; cysteine (C) carbamidomethylation], [Variable Modification; methionine (M) oxidation/ N-terminal acetylation/ serin (S) threonine (T) tyrosine (Y) phosphorylation], [Maximum Variable Modifications; 3]*. Peptide validation was performed using the Percolator algorithm, and only high confidence peptides (false discovery rate (FDR) < 1%) were used for protein inference and MS1-based LFQ. For phosphoproteomic analysis, the site localization probability of phosphorylated amino acid was calculated using IMP-ptmRS node implemented in Proteome Discoverer software.

To calculate abundance fold change for different biological conditions, the phosphopeptides with more than or equal to three quantitative values in each sample group, were selected (the number of 14,645 peptide groups in Supplemental Table 2). The abundances were normalized by Levenberg-Marquardt optimization ^61^ and missing values were replaced by the minimum value. The source code was described in a previous study ^44^.

### Analysis of proteomic data

Unless otherwise noted, basic calculations related to proteomic data were performed in Excel program. For visualization of phosphopeptide intensities as a heat map, mean values for each peptide were scaled to 100 (Supplemental Table 6). GO enrichment analysis was performed as follows: the protein IDs in each Excel sheet (No.1, 4, 7 in Supplemental Table 2) were uploaded to PANTHR (https://www.pantherdb.org/), and the background database was set as Arabidopsis all gene IDs. The GO terms shown in the Figures were selected as GO terms with less than 1% FDR.

### Measurement of ABA-induced ROS production in guard cells

Measurement of ROS levels in guard cells was performed according to a previous study ^54^. Briefly, dark adapted epidermal strips were incubated in stomatal opening buffer [5 mM MES-bistrispropane (pH 6.5), 50 mM KCl, and 0.1 mM CaCl_2_] for 1.5 h in the dark or in red light (RL) [50 µmol m^−2^ s^−1^] plus blue light (BL) [10 µmol m^−2^ s^−1^] at 24°C and then treated with 50 µM 2ʹ,7ʹ-dichlorofluorescin diacetate (H_2_DCFDA) for 30 min. After washing the epidermis to remove excess dye, ABA-induced ROS production was measured by adding 10 µM ABA to under the RL plus BL and the epidermis was incubated for another 30 min. DCF fluorescence was observed using a fluorescence microscopy (Eclipse TS100; Nikon), and the fluorescence intensity was measured using ImageJ software.

### Whole-cell patch-clamp measurement of Arabidopsis GCPs

*I_Ca_* channel currents in Arabidopsis GCPs were recorded according to previous studies ^27,62^. Whole-cell currents were recorded using a MultiClamp 700B patch-clamp amplifier (Molecular Devices, San Jose, CA) and the data were analyzed with pCLAMP 10.3 software (Molecular Devices). The pipette solution contained 10 mM BaCl_2_, 0.1 mM DTT, 5 mM NADPH, 4 mM EGTA, and 10 mM HEPES-Tris (pH 7.1). The bath solution contained 100 mM BaCl_2_, 0.1 mM DTT and 10 mM MES-Tris (pH 5.6). The osmolarity of the solutions was adjusted with D-sorbitol to 500 mmol kg^−1^ (pipette) and 485 mmol kg^−1^ (bath).

### Accession numbers

Sequence data for the genes described in this article are avairable at TAIR (http://www.arabidopsis.org/) under the following accession numbers: MAP4K1/AtMAP4Kα1 (AT1G53165), MAP4K2/AtMAP4Kα2 (AT3G15220), MAP4K3/SIK1 (AT1G69220), MAP4K4/TOT3 (AT5G14720), MAP4K5/TOI5 (AT4G24100), MAP4K6/TOI4 (AT4G10730), MAP4K7/TOI3 (AT1G70430), MAP4K8/TOI1 (AT1G79640), MAP4K9/TOI2 (AT1G23700), MAP4K10/BLUS1 (AT4G14480), SLAC1 (AT1G12480).

## Data availability

LC-MS/MS raw data and of phosphoproteomic/proteomic analysis have been deposited in Japan Proteome Standard Repository/ Database (jPOSTP) Proteomics; https://repository.jpostdb.org/preview/28177253465b9ac41e6a1b, Access key 6697/Phosphoproteomics; https://repository.jpostdb.org/preview/26082790465b9b5348a2b5, Access key 3128).

## Acknowledgments

We thank Dr. Tsuyoshi Nakagawa (Shimane University, Japan) for providing the R4pGWB 501 vector. We are grateful to the ABRC for providing Arabidopsis T-DNA insertion mutants. We thank Mrs. Sakurako Hosotani for her experimental assistance. This work was supported in part by the Japan Society for the Promotion of Science (JSPS) KAKENHI grants JPMJPR13B3, JP25840103, JP19H03240, JP21H05654 to T.U., JP21J10962 to Y.K., Moonshot program 20350427 to T.U.

## Author contributions

K.Y. and T.U. designed experiments and wrote the manuscript with contributions from all authors. K.Y., S.K., H.T., Y.L., A.O., Y.K., S.Y. and I.C.M. performed experiments. A.T. provided methods and critically advised experiments.

## Conflict of interest

The authors declare no conflict of interest.

---

**Fig. S1.**
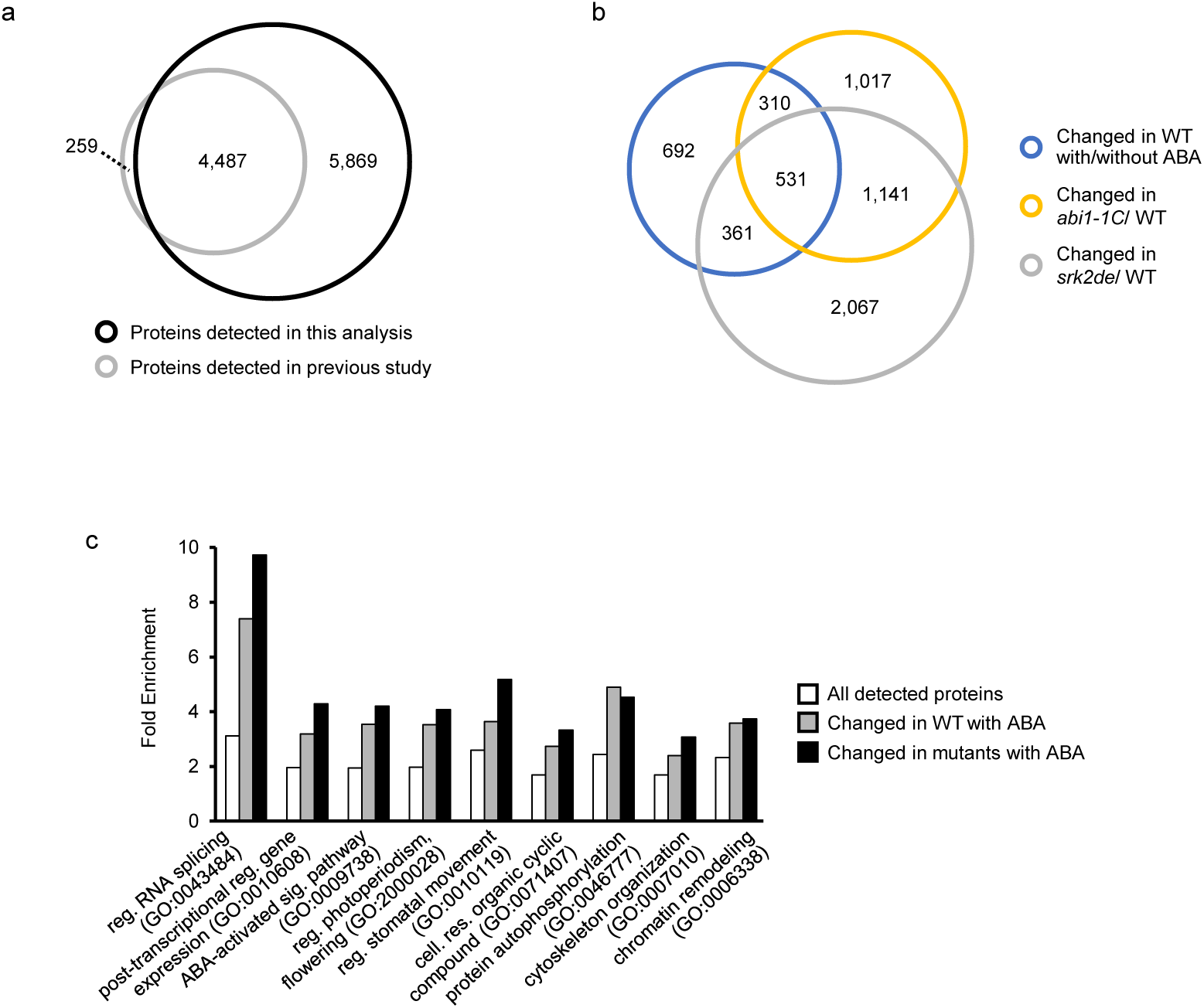
An overview of phosphoproteomic analysis using Arabidopsis guard cells. **a,** Venn diagram shows the overlap between the proteins detected in this study and a previous study ^22^ on Arabidopsis GCPs. **b,** Venn diagram shows the overlap between ABA-responsive phosphopeptides in Col-0 wild-type (WT) and up/down-regulated phosphopeptides in *srk2de* or *abi1-1C*. **c,** Gene Ontology (GO) enrichment analysis. All of detected proteins or proteins with significant changes in ABA or mutants were assigned. Enriched GO terms were evaluated by PANTHR program and GO terms with less than 1% FDR were employed.

**Fig. S2.**
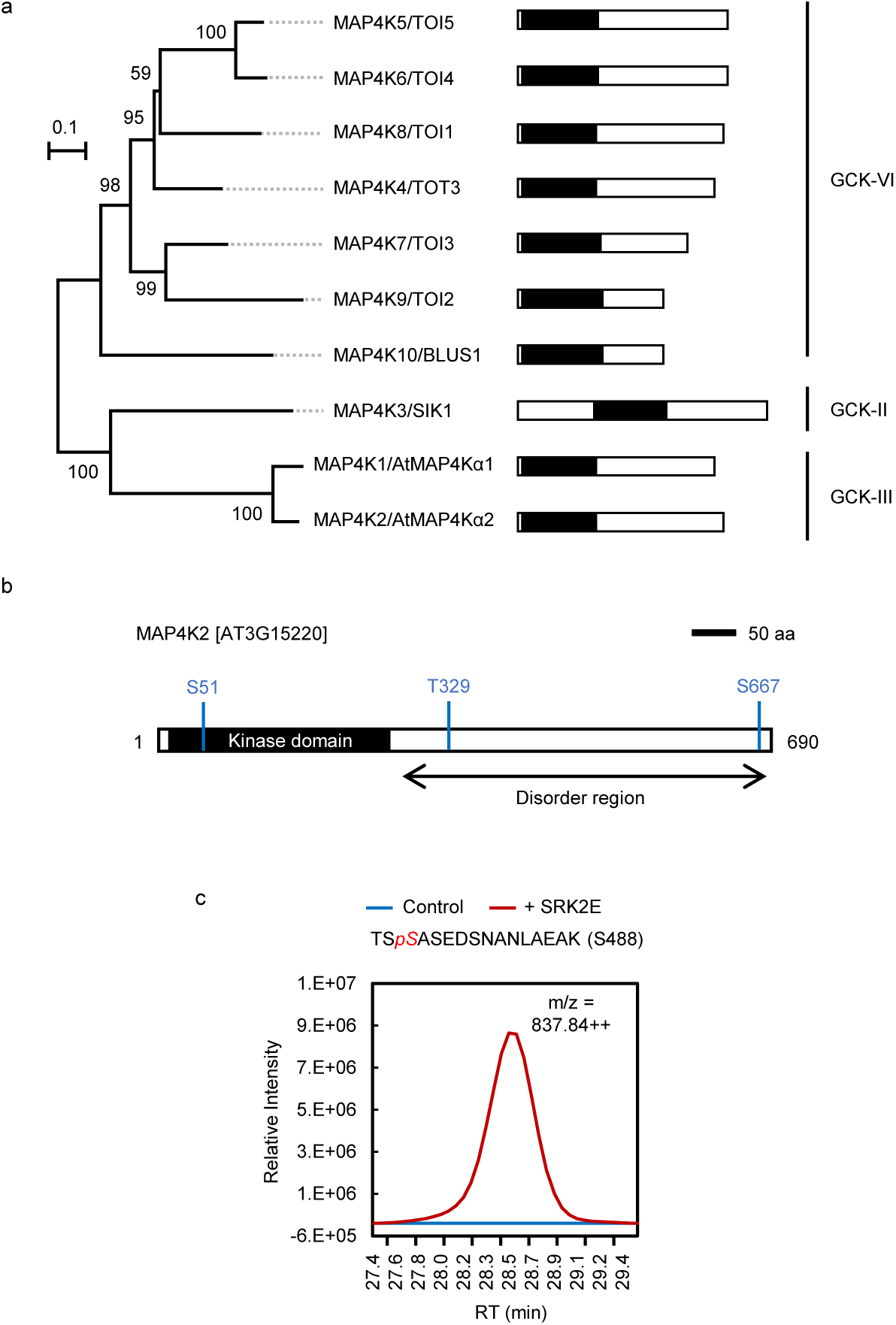
MAP4K2 is directly phosphorylated by SRK2E *in vitro*. **a**, A phylogenetic tree of MAP4K gene family in Arabidopsis *thaliana*. A schematic of the protein sequences is shown in right, including the position of kinase domain (black) and red line indicates conserved Ser or Thr (S/T) for MAP4K1 Ser-479. The evolutionary history was inferred using the Neighbor-Joining method. The percentage of replicate trees in which the associated taxa clustered together in the bootstrap test (1000 replicates) are shown next to the branches. All positions containing gaps and missing data were eliminated. All protein sequences were directly obtained from the Araport11. The phylogenic analysis was performed using the full length of each protein. Arabidopsis MAP4K family can be classified to germinal center kinase (GCK)-type II, III or VI based on classifications in mammalian. **b,** A schematic of Arabidopsis MAP4K2. Phosphorylation sites identified in the phosphoproteomic analysis was shown in blue line. MAP4K2 have kinase domain (black) in N-terminus and long disorder region in C-terminus. **c,** *In vitro* phosphorylation assay using an active MBP-SRK2E incubated with kinase-dead form of MBP-MAP4K2 (MAP4K2 K44N) followed by LC-MS/MS analysis. LC-MS/MS analysis detected the extracted ion chromatogram (XIC) of MAP4K2 peptide [TS*pS*ASEDSNANLAEAK] containing phosphorylated Ser-488 only when incubated with SRK2E, but not when incubated alone (Control).

**Fig. S3.**
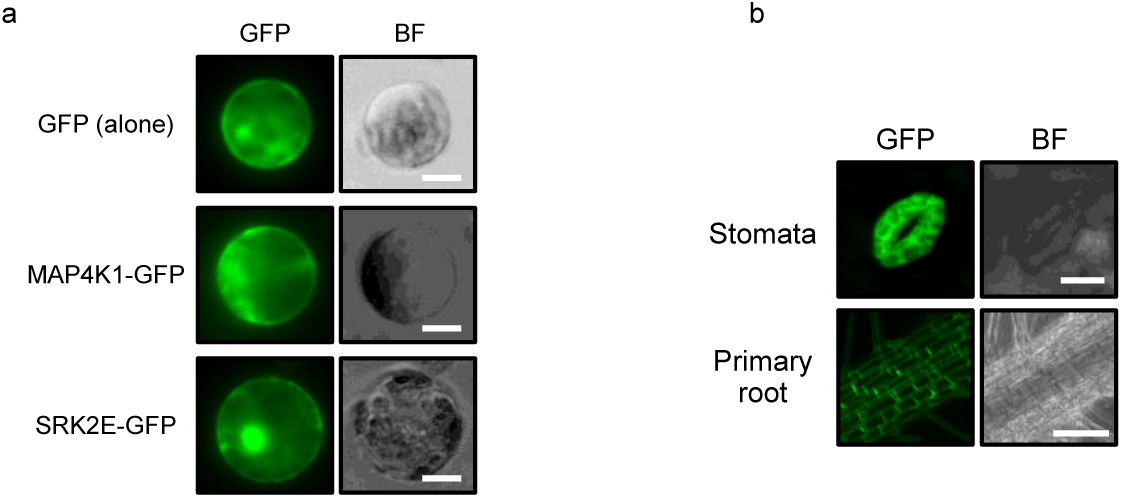
Subcellular localization analysis of MAP4K1-GFP protein. **a,** Subcellular localization of GFP (alone), MAP4K1-GFP and SRK2E-GFP in Arabidopsis mesophyll cell protoplasts (MCPs). Scale bars indicate 25 µm. **b,** Subcellular localization MAP4K1-GFP in stomata or primary root of *35S:MAP4K1-GFP* Arabidopsis transgenic plants. Scale bars indicate 15 µm or 100 µm for the images of stomata or primary root, respectively.

**Fig. S4.**
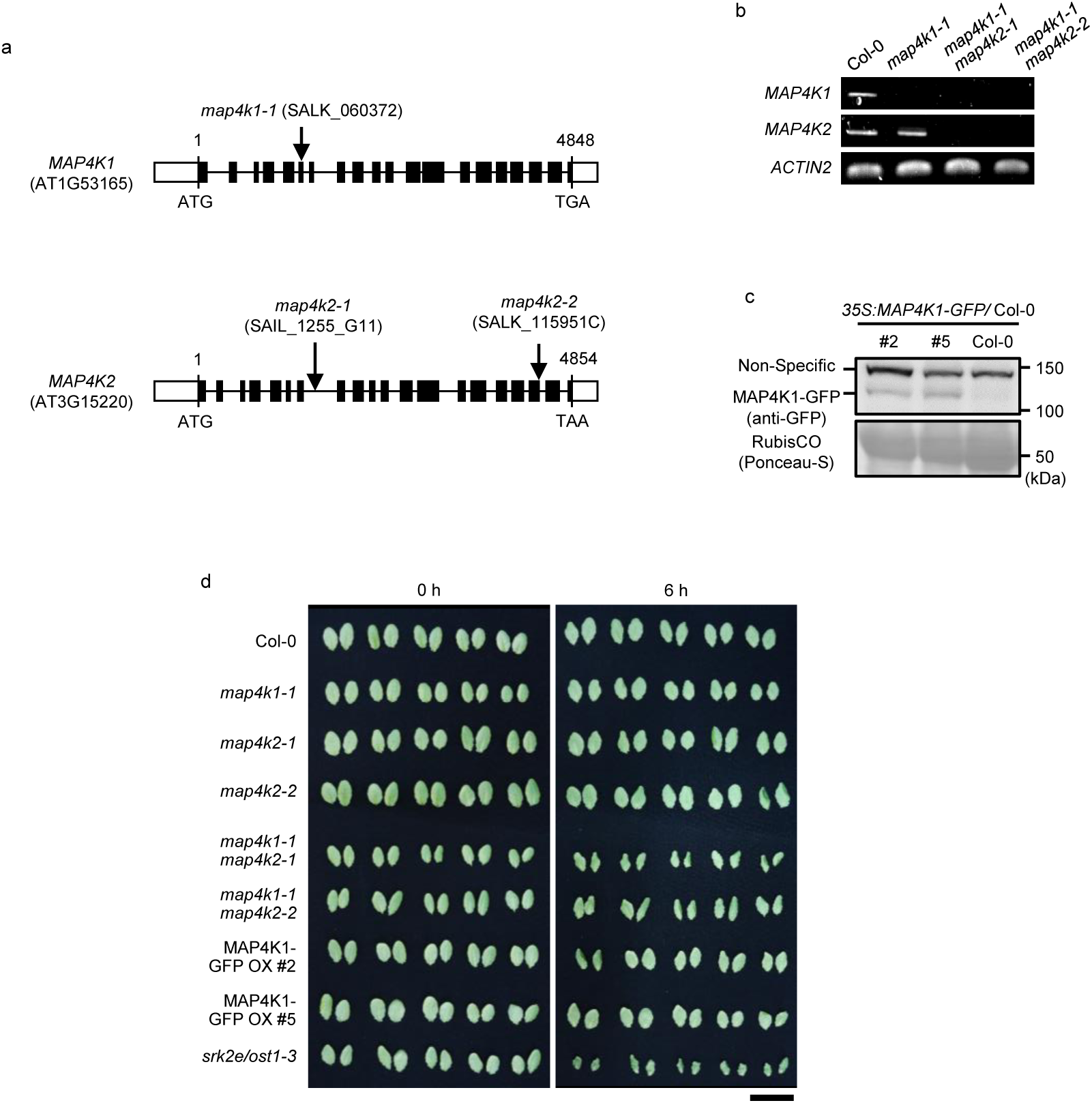
Established Arabidopsis MAP4K1/2 KO mutants and MAP4K1-GFP OX plants. **a,** Schematic representation of the genomic DNA sequence shows T-DNA insertions in map4k1 or map4k2-1/-2. Solid boxes and lines indicate exon and intron, respectively. **b,** RT-PCR analysis of Col-0 and *map4k1*, *map4k1map4k2-1* and *map4k1map4k2-2. ACTIN2* was used as a positive control. **c,** Western blot analysis of *35S:MAP4K1-GFP/ Col-0*. MAP4K1-GFP was detected with an anti-GFP antibody, and RubisCO was indicated by CBB staining. **d,** Images show the representative detached rosette leaves from four-week-old plants of Col-0, *srk2e/ost1-3*, *MAP4K1/2* knockout mutants and *MAP4K1-GFP OX* plants. Photos were taken at 0h or 6h after detached. Scale bar indicates 3 cm.

**Fig. S5.**
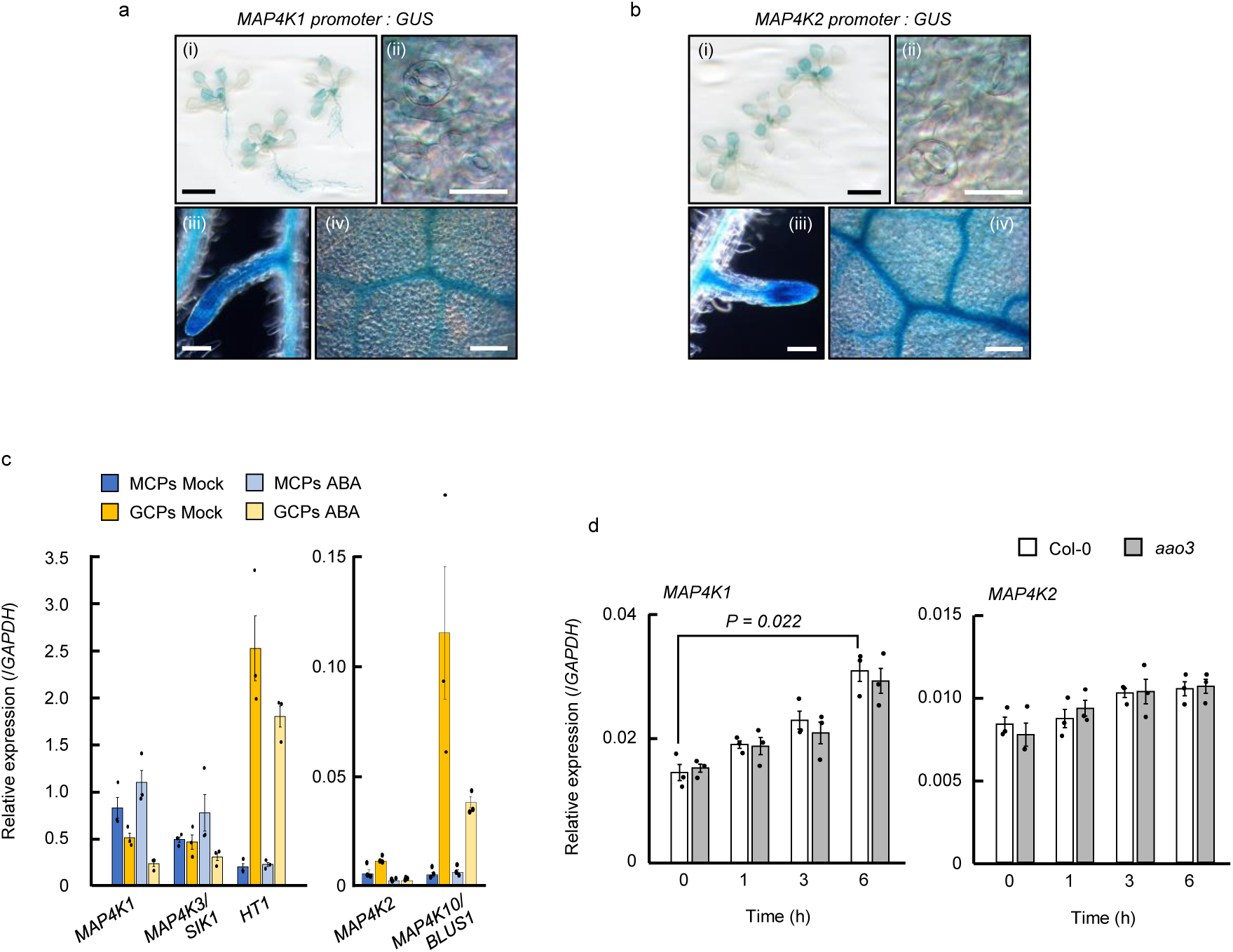
Tissue specific expression pattern of *MAP4K1* and *MAP4K2*. **a,b,** Histochemical GUS staining of 2-weeks-old *MAP4K1 or MAP4K2 promoter : GUS* plants. Images show the whole seedlings (i), stomata (ii), lateral root (iii) and vein (iv), respectively. Scale bars show 1 cm (i), 25 µm (ii) and 100 µm (iii and iv), respectively. **c,** Relative gene expression level of *MAP4K1*, *MAP4K2*, *MAP4K3/SIK1*, *MAP4K10/BLUS1* in guard cell protoplasts (GCPs) or mesophyll cell protoplasts (MCPs). HT1, a guard cell specific protein kinase, is a marker gene for guard cell. **d**, Osmotic stress-induced relative expression level of *MAP4K1* and *MAP4K2* in 2-weeks-old Arabidopsis Col-0 and *aao3,* which mutant defects ABA biosynthesis, seedlings treated with 400 mM sorbitol for indicated periods. *GAPDH* was used as an internal control. Bars indicate means ± SE (n = 3), and significance was determined using two-tailed Student’s t-test adjusted with Bonferroni.

**Fig. S6.**
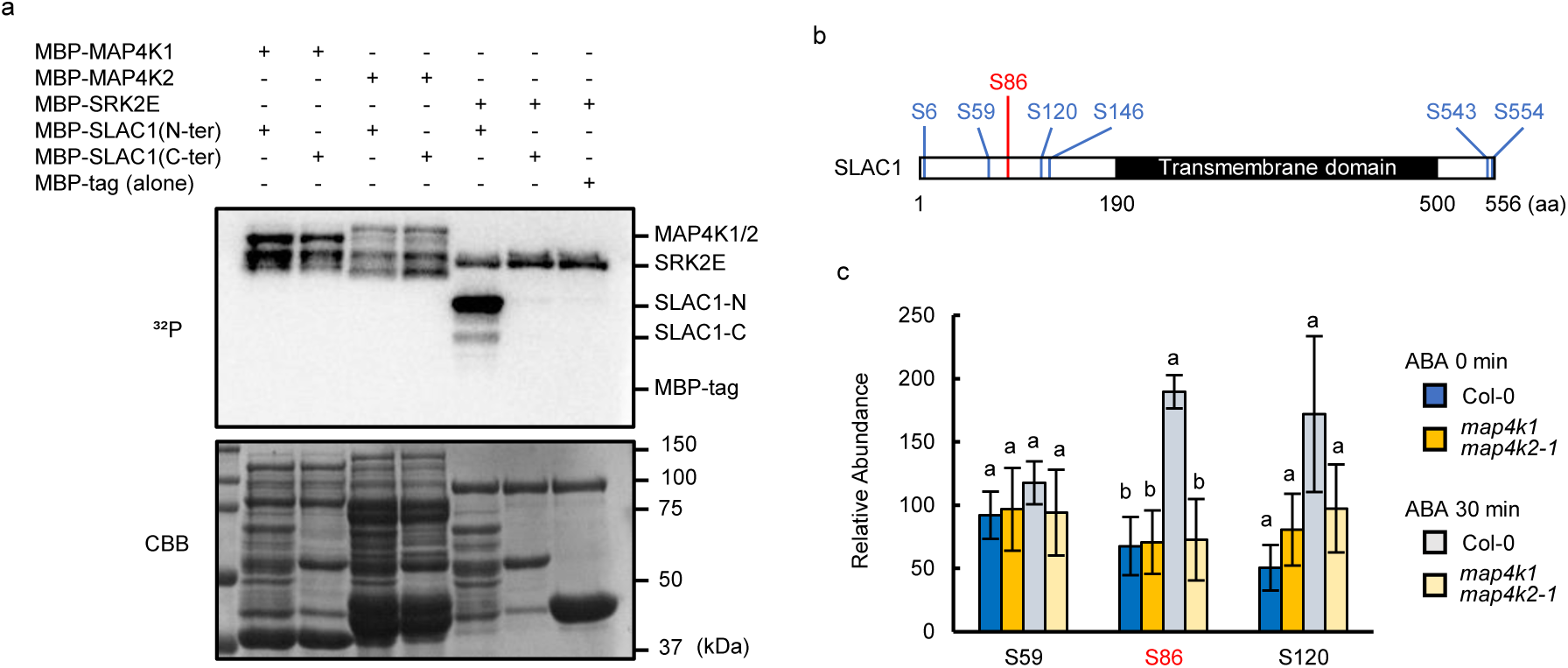
MAP4K1 and MAP4K2 can not phosphorylate N-terminus and C-terminus of SLAC1, but may contribute to the phosphorylation of SLAC1 Ser-86 *in planta*. **a**, *In vitro* phosphorylation assay was performed using SLAC1 fragments of N-terminus (1-189 aa) or C-terminus (501-556 aa) as substrates in the presence of MBP-tagged SRK2E, MAP4K1 or MAP4K2. Phosphorylation levels were detected by autoradiography. CBB staining shows protein loading in each lane. **b**, Phosphorylated sites of SLAC1 protein are identified in a phosphoproteomic analysis with GCPs of Col-0 and *map4k1map4k2-1*. Phosphopeptide samples were prepared from Col-0 or *map4k1map4k2-1* GCPs with/without 10 µM ABA for 30 min. **c**, Relative abundance of the phosphopeptide included Ser-59, 86 and 120 of SLAC1 in Col-0 or *map4k1map4k2-1* GCPs. The mean intensities were scaled to the same range for each phosphopeptide. Data represent means ± SE (n = 4, biologically independent replicate). Different letters indicate significant differences (Tukey’s test, P < 0.05).

**Fig. S7.**
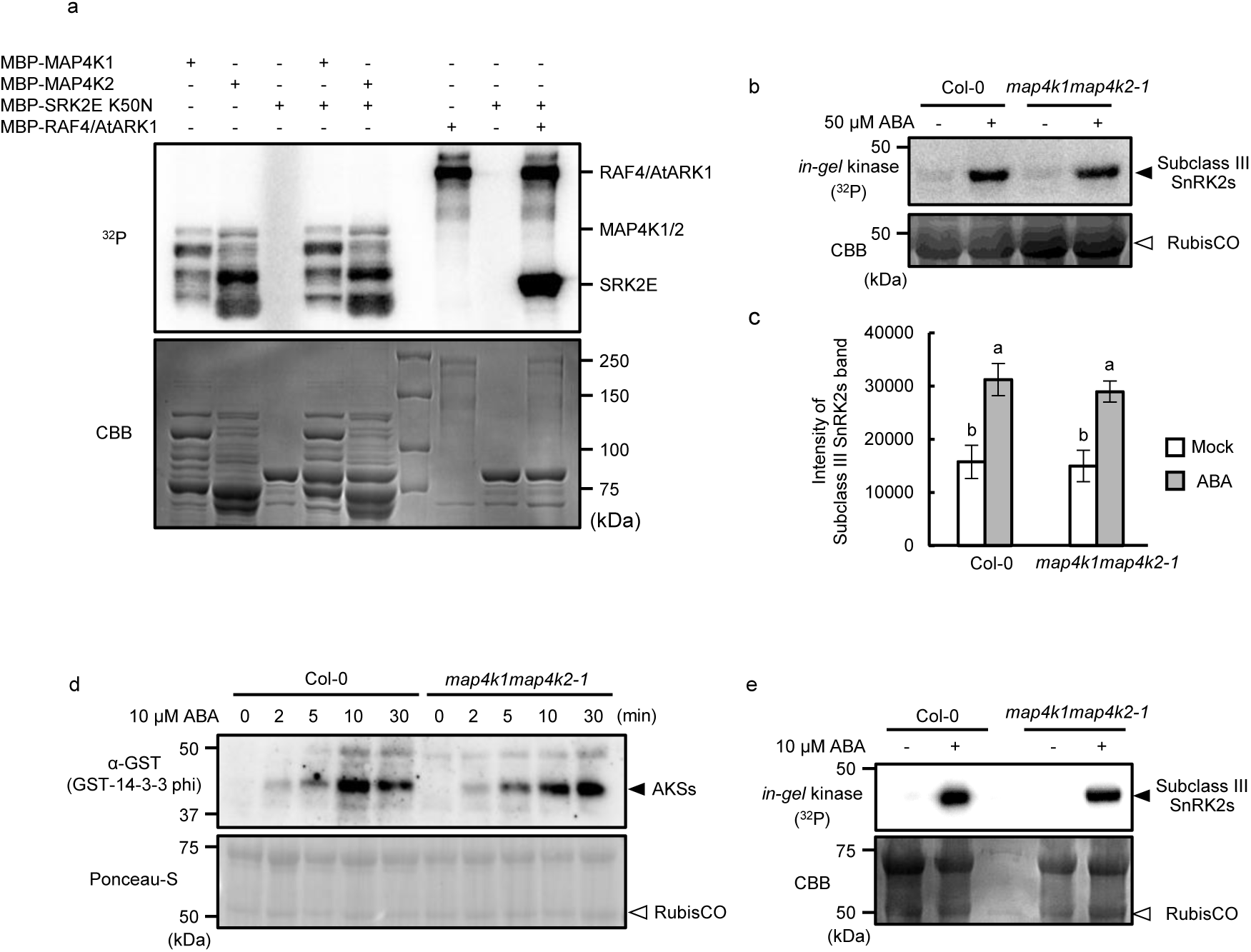
MAP4K1/2 is not involved in ABA-dependent activation of SnRK2s. **a,** *In vitro* phosphorylation assay was performed using MBP-SRK2E^K50N^ as a substrate. MBP-tagged MAP4K1, MAP4K2 or RAF4/AtARK1 were incubated with MBP-SRK2E^K50N^ in the presence of [γ-^32^P]ATP. Phosphorylation levels were detected by autoradiography. CBB-staining shows protein loading in each lane. **b,** *in-gel* phosphorylation assay of protein extracts from two-week-old seedlings of Col-0 and *map4k1map4k2-1* treated with 50 µM ABA for 30min. Histone III-S was used as a substrate. Black and open arrows indicate SnRK2s and RubisCO, respectively. Autoradiography (^32^P) and coomassie brilliant blue (CBB) staining show phosphorylation and protein loading, respectively. **c,** Quantification of SnRK2s in (b). SnRK2s were quantified using ImageJ software. Data are presented as means ± SE (n = 4). Different letters in indicate significant differences (Tukey’s test, P < 0.05). **d,** Far-western blot analysis of 14-3-3 protein using protein extracts from GCPs of Col-0 and *map4k1map4k2-1* treated with 10 µM ABA. Black and open arrows indicate AKSs and RubisCO. **e,** In-gel phosphorylation assay of protein extracts from GCPs of Col-0 and *map4k1map4k2-1* treated with 10 µM ABA for 10 min. Histone III-S was used as a substrate. Black and open arrows indicate SnRK2s and RubisCO, respectively. Autoradiography (^32^P) and coomassie brilliant blue (CBB) staining show phosphorylation and protein loading, respectively.

**Fig. S8.**
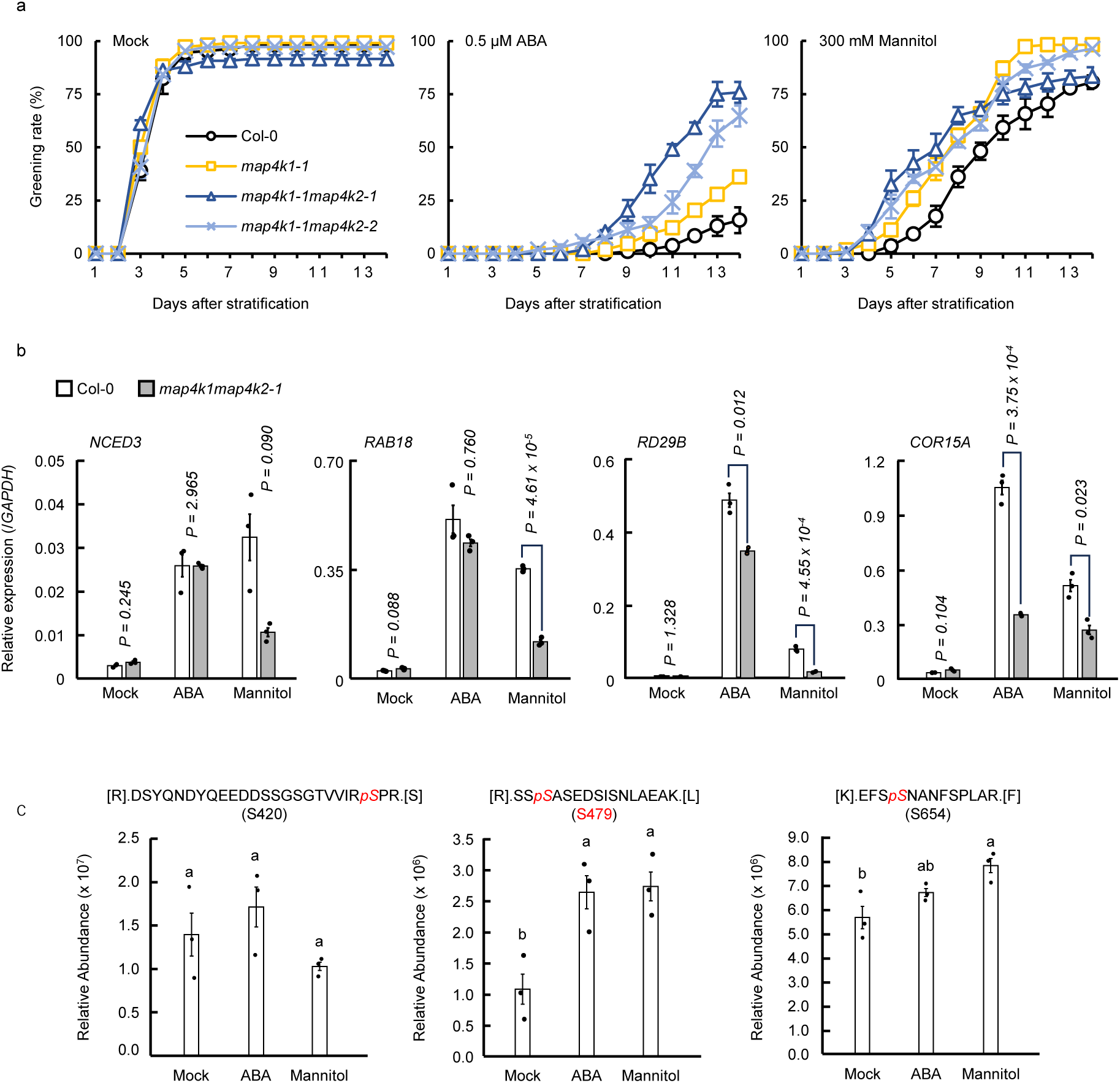
Phosphorylation of Ser-479 at MAP4K1 may not affect autophosphorylation *in vitro*. *In vitro* phosphorylation assay using an active MBP-MAP4K1 WT/S479A/S479D or MBP-MAP4K2 WT/S488A/S488D. MAP4K1 or MAP4K2 were incubated in the presence of [γ-^32^P]ATP. Autophosphorylation levels were detected by autoradiography. CBB staining showed protein loading in each lane.

**Fig. S9.**
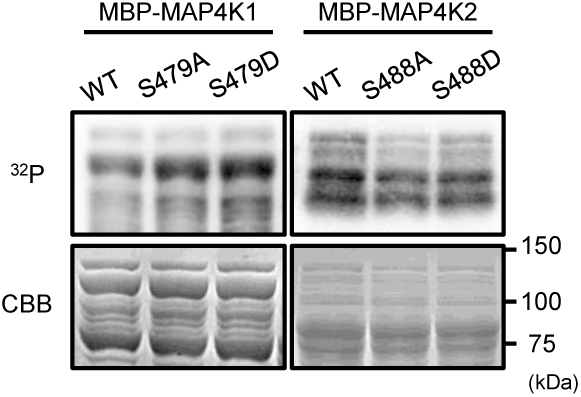
MAP4K1/2 is involved in ABA- and osmotic stress-response in Arabidopsis seedlings. **a,** Quantification of the cotyledon greening rates of Col-0 wild-type, *map4k1-1*, *map4k1-1map4k2-1* and *map4k1-1map4k2-2* on GM agar medium with or without 0.5 µM ABA/ 300 mM Mannitol. Data presents means ± standard error (n=3). Each replicate contains 36 seeds. **b,** Relative expression levels of ABA- or osmotic stress-responsive genes in 2-week-old Arabidopsis Col-0 and *map4k1-1map4k2-1* seedlings treated with 50 μM ABA or 400 mM mannitol for 3 h. *GAPDH* was used as an internal control. Bars indicate means ± SE (n = 3), and significance was determined using two-tailed Student’s t-test adjusted with Bonferroni. **c**, Relative abundance of MAP4K1 peptides including phosphorylated Ser-420, Ser-479 or Ser-654. Peptide samples were prepared from 2-weeks-old Arabidopsis seedling with/without 50 µM ABA or 400 mM Mannitol for 30 min. Data bars represent means ± SE (n = 3, biologically independent replicate). Different letters indicate significant differences (Tukey’s test, P < 0.05).

**Fig. S10.**
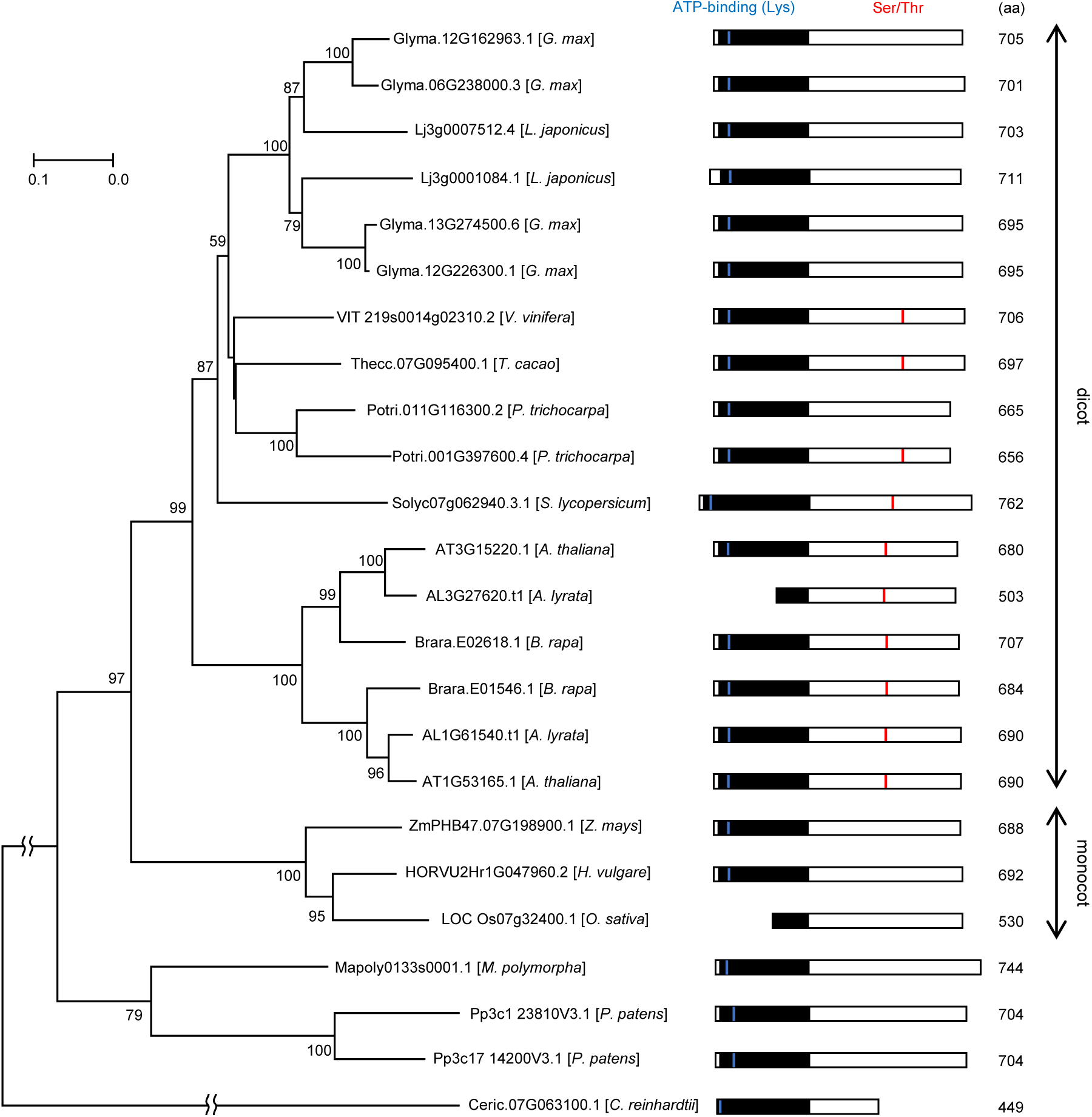
A phylogenetic tree of *MAP4K1* orthologs and its homolog pair in the green lineage. The evolutionary history was inferred using the Neighbor-Joining method. The percentage of replicate trees in which the associated taxa clustered together in the bootstrap test (1000 replicates) are shown next to the branches. All positions containing gaps and missing data were eliminated. All protein sequences were directly obtained from the Phytozome v13. The phylogenic analysis was performed using the full length of each protein. A schematic of the protein sequences is shown in right, including the position of kinase domain (black) and red line indicates conserved Ser or Thr (Ser/Thr) for *A. thaliana* MAP4K1 Ser-479, which is identified in the phosphoproteomic analysis with GCPs as an ABA-responsive phosphorylation site. ATP-binding site (Lys) is shown in the bule line and the numbers in right of each protein schematic shows the amino acid length, respectively.

## Notes

### Competing Interest Statement

The authors have declared no competing interest.

